# Diet-conditioned microbiota enhances fecal microbiota transplantation efficacy in alcoholic liver disease through caproic acid-PPARα signaling

**DOI:** 10.64898/2026.03.25.714243

**Authors:** Nishu Choudhary, Ashi Mittal, Sandeep Kumar, Kavita Yadav, Anupama Kumari, Deepanshu Maheshwari, Pragyan Parimita Rath, Jaswinder Singh Maras, Asif Mohmmed, Anupam Kumar, Shiv K Sarin, Shvetank Sharma

## Abstract

**Background and Aim:** Fecal microbiota transplantation (FMT) in Alcohol-related liver disease (ALD) has shown therapeutic potential, with variable efficacy and unclear mechanism. Because dietary protein influences gut microbiota composition, we hypothesized that donor dietary preconditioning could enhance FMT efficacy. We therefore examined in a murine ALD model if high-protein donor diet improves FMT outcome.

**Methods:** ALD was induced in C57BL/6N mice using a Lieber-DeCarli ethanol diet combined with thioacetamide administration for 12 weeks. FMT was performed using stool from diet-modulated donors, and recovery was assessed on day7 post-FMT. Multi-omics analysis using 16s rRNA and mass spectroscopy was performed for Gut microbiota composition, plasma- and stool-metabolome, and hepatic proteomes. Multi-omics outcomes were validated in ALD animal and Huh7 hepatocytes.

**Results:** Both protein-based FMTs improved ALD recovery; Veg-FMT indicated superior efficacy, significantly reducing hepatic injury (AST 1.2FC, p=0.002; bilirubin 1.2FC, p=0.03; steatosis 1.7FC,p=0.01) and restoring gut barrier integrity (occludin 1.5FC,p=0.04; mucin 2 2.2FC, p=002; and plasma endotoxin 1.7FC, p=0.02). A significant 2FC increase was observed in *Lachnospiraceae* NK4A136, *Coriobacteriaceae* UCG-002, and short-chain fatty acids, particularly caproic acid. Functional validation confirmed that caproic acid promoted hepatic fatty acid β-oxidation through PPARα-dependent mechanisms, reducing triglyceride accumulation and lipogenesis in both cellular and animal models.

**Conclusion:** Donor preconditioning with a plant-protein enriched diet enhances FMT efficacy in ALD by gut microbiota modulation with increased metabolites like caproic acid. These findings highlight a microbiota-metabolite-host axis through which diet-modulated FMT improves hepatic lipid metabolism and injury, and identifies a pathway via which FMT imparts its effect.

**Significance:** This study identifies a mechanistic basis for improving fecal microbiota transplantation (FMT) efficacy in alcohol-related liver disease (ALD) by demonstrating that dietary preconditioning of donor microbiota improves therapeutic outcomes. We show that plant protein–modulated donor microbiota supplements abstinence-associated recovery through increased production of the microbial associated metabolite caproic acid, which promotes hepatic fatty acid β-oxidation via PPARα signaling. These findings highlight donor dietary conditioning and microbiota-derived metabolites, rather than microbial composition alone, as important determinants of FMT efficacy. The results suggest that microbial associated metabolites such as caproic acid may represent potential therapeutic targets or biomarkers to enhance and standardize microbiota-based interventions in ALD. Although the current work is based on a murine model, the identified microbiota–metabolite–host metabolic axis provides a framework for future translational studies aimed at optimizing FMT strategies in liver disease.

## INTRODUCTION

Among alcohol-associated liver disease (ALD) patients, a majority develop steatosis, with only 10–35% developing hepatitis, of which 8–20% eventually progress to cirrhosis (1). The pathophysiology of ALD is influenced by compositional shifts in intestinal bacteria (2,3). Alcohol intake reduces bacterial diversity and increases the abundance of *Proteobacteria* and *Enterobacteria* and decreases the abundance of *Bacteroidetes* and Firmicutes, specifically *Lactobacillus* species, leading to intestinal barrier dysfunction and endotoxemia (3–5).

Restoration of gut microbial homeostasis has therefore emerged as a promising therapeutic strategy. Fecal microbiota transplantation (FMT) has indicated beneficial effects in experimental models and clinical studies of alcoholic hepatitis, including improvements in liver injury and survival (5–9). However, responses to FMT remain variable, probably due to the condition of the patient or an inept donor bacterial composition (10). These findings suggest that donor microbiota characteristics may critically influence therapeutic efficacy, highlighting the need to better understand factors that shape donor microbial function.

Diet is one of the strongest determinants of gut microbiota composition and metabolic activity (11,12). Short-term dietary changes can rapidly alter microbial communities and their metabolite production. Although dietary fiber is widely recognized as a major substrate for microbial fermentation, dietary proteins that escape digestion in the small intestine also reach the colon and can serve as substrates for microbial metabolism (13–15). Microbial fermentation of amino acids produces a variety of metabolites capable of modulating host metabolic and inflammatory pathways (16,17).

Despite these associations, the influence of donor dietary conditioning on FMT efficacy in ALD remains poorly understood. In this study, we investigated whether preconditioning donor microbiota with diets enriched in vegetable- or egg-derived proteins could enhance FMT-mediated recovery in a murine model of ALD. Using integrated microbiome, metabolomic, and proteomic analyses, we suggest that vegetable protein–modulated microbiota significantly improve recovery from ALD. This effect is associated with enrichment of short-chain fatty acid producing bacteria and increased production of caproic acid, a microbial associated metabolite that activates hepatic PPARα signaling to enhance fatty acid β-oxidation and reduce hepatic lipid accumulation. These findings highlight dietary modulation of donor microbiota as a potential strategy to improve microbiota-based therapies in alcohol-related liver disease.

## RESULTS

### Protein-conditioned FMT enhances recovery from alcohol-induced liver injury

To determine whether donor dietary conditioning improves the therapeutic efficacy of fecal microbiota transplantation (FMT), donor mice were preconditioned with either standard diet, vegetable protein diet, or egg protein diet prior to microbiota transfer (Fig. 1A). FMT was administered to ALD mice following abstinence, and liver injury parameters were assessed seven days after transplantation.

**Figure 1:**
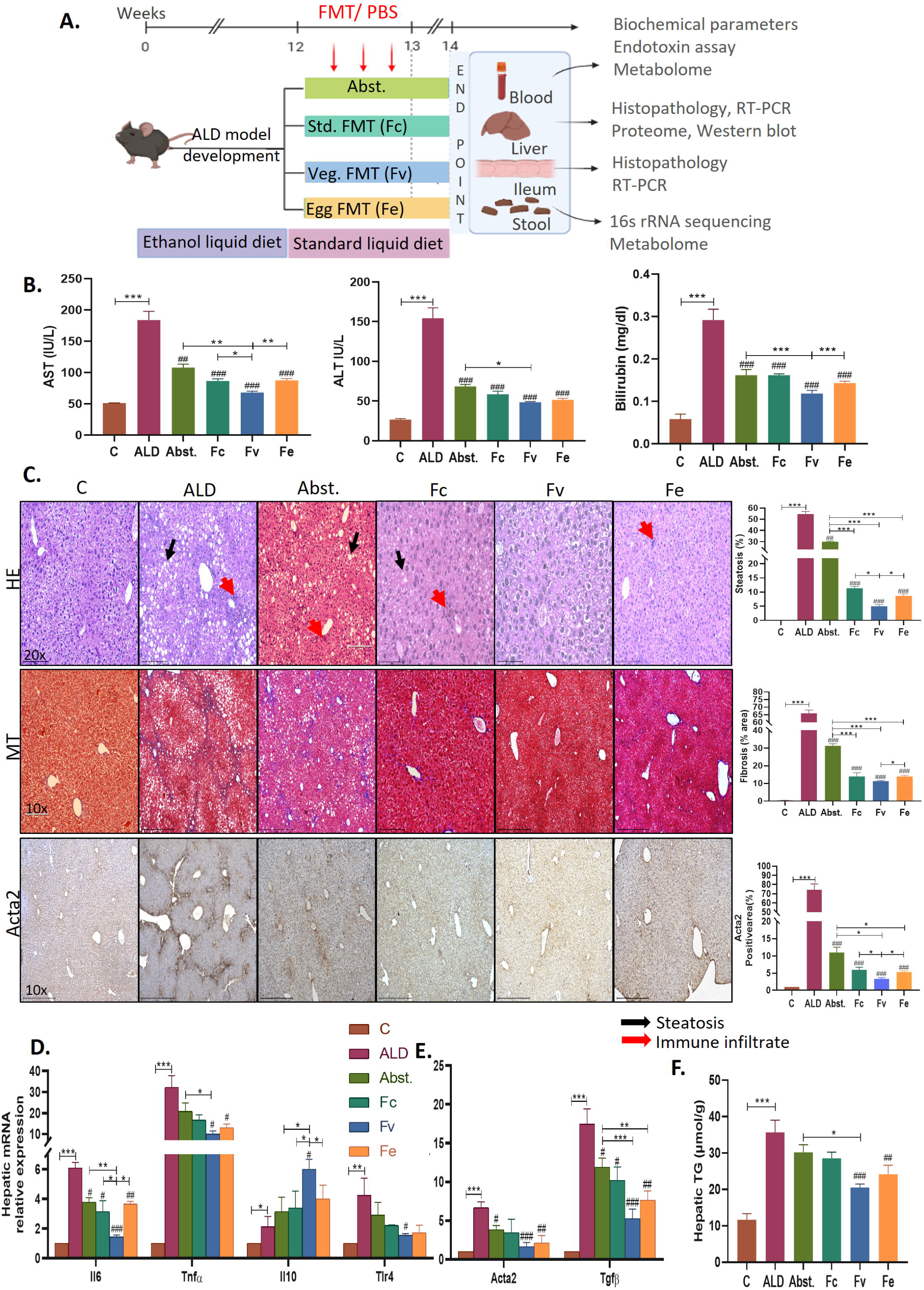
Protein-conditioned FMT improves liver function. **(A)** Study design- In separate setups, FMT was given to ALD mice (3 alternate days) from donors fed one of three diets—a standard diet, a vegetable protein diet or an egg protein diet—followed by collection of tissues and blood seven days after FMT. **(B)** Overall, FMT reduced serum AST, ALT and bilirubin levels, with an enhanced reduction in Veg-FMT (all p<0.001). **(C)** Representative micrographs of liver histology by hematoxylin and eosin (H&E) and Masson’s trichome (MT) staining and immunohistochemistry for smooth muscle actin protein (α-sma) showing decreased hepatic steatosis (p<0.001) and fibrosis-associated staining (MT and α-sma positive area, p=0.01in both) in protein-conditioned FMT with statistical analysis. **(D)** Compared with egg-FMT, Veg-FMT led to greater reductions in hepatic Il6 (p=0.04), Tnfα (p=0.06), Tlr4 (p=0.05), **(E)** Acta2 (p=0.001) and Tgfβ(p<0.001) and increased IL10 (0.02) mRNA expression. **(F)** Hepatic TGs were significantly reduced in protein-conditioned FMT (Veg-FMT: 1.74FC, p<0.001; Egg-FMT: 1.47FC,p=0.009). The data are presented as the means ± SEMs (standard error of the mean). *p < 0.05, **p < 0.01, ***p < 0.001 represent intergroup statistics; #p < 0.05, ##p < 0.01, ###p < 0.001 represent the ALD group (one-way ANOVA followed by Tukey’s multiple comparison test).

Abstinence alone resulted in partial recovery from liver injury, as indicated by increased food intake (fold change >2, p<0.05) and a reduction in the liver-to-body weight ratio compared with ALD mice (Additional file 1, S. Fig. 1B-C). Serum biochemical analysis confirmed successful induction of liver injury in ALD mice, with significantly elevated levels of AST, ALT, and bilirubin compared with controls. Abstinence significantly reduced these markers relative to ALD mice (Fig. 1B).

FMT further improved liver injury beyond abstinence alone. Standard FMT significantly reduced AST (2.1FC), ALT (2.6FC), and bilirubin (1.8FC) compared with ALD mice (all p<0.001). Protein-conditioned FMT produced greater improvement, with Veg-FMT showing the most pronounced effect. Veg-FMT reduced AST by 2.7FC, ALT by 3.2FC, and bilirubin by 2.6FC compared with ALD mice (all p<0.001) (Fig. 1B).

Histological examination supported these findings. Hematoxylin-eosin staining revealed marked hepatic steatosis and inflammatory infiltration in ALD mice, which were partially improved after abstinence and further ameliorated following FMT treatment (Fig. 1C). Veg-FMT produced the greatest reduction in steatosis, showing an approximately 11FC decrease compared with ALD mice (p<0.001).

Consistent with the histological observations, hepatic triglyceride levels were significantly reduced following protein-modulated FMT. Veg-FMT reduced hepatic triglycerides by 1.74FC (p<0.001), while Egg-FMT reduced triglycerides by 1.47FC (p=0.009) relative to ALD mice (Fig. 1F).

Veg-FMT also significantly improved inflammatory and fibrosis-associated markers. Expression of pro-inflammatory cytokines Il6 (2.54FC reduction, p=0.04), Tnfα (1.29FC reduction, p=0.06), and Tlr4 (1.11FC reduction, p=0.05) decreased following Veg-FMT, while the anti-inflammatory cytokine Il10 increased 1.5FC (p=0.02) (Fig. 1D). Similarly, fibrogenic genes Acta2 (4.1FC reduction, p<0.001) and Tgfβ (3.3FC reduction, p<0.001) were significantly downregulated (Fig. 1E).

Collectively, these findings indicated that dietary conditioning of donor microbiota enhances early recovery from alcohol-induced liver injury, with vegetable protein-modulated microbiota showing the strongest protective effect.

### Veg-FMT restores intestinal barrier integrity and reduces endotoxemia

Chronic alcohol exposure disrupts intestinal barrier integrity, leading to increased gut permeability and translocation of microbial products such as endotoxin, which contribute to hepatic inflammation through the gut–liver axis. To determine whether protein-conditioned fecal microbiota transplantation (FMT) improves intestinal barrier function, we examined intestinal histology, tight-junction protein expression, antimicrobial peptides, inflammatory markers, and circulating endotoxin levels following treatment.

Histological analysis of intestinal tissues revealed marked epithelial damage and shortening of intestinal villi in ALD mice compared with control animals (Fig. 2A). Although abstinence resulted in partial recovery of intestinal morphology, FMT treatment further improved epithelial structure. Notably, Veg-FMT restored villus architecture to near-normal morphology, whereas Egg-FMT and standard FMT showed only moderate improvement (Additional file 1, S. Fig. 2A). Consistent with these histological findings, ALD mice exhibited significantly reduced expression of key tight-junction proteins, indicating impaired intestinal barrier function. Veg-FMT significantly increased mRNA expression of Zo-1 (2.04FC, p = 0.05), Occludin (2.31FC, p = 0.03), and Claudin-3 (1.63FC, p = 0.01), and significantly increased ZO-1 protein expression compared with ALD mice (Fig. 2A). These results suggest that Veg-FMT effectively restores epithelial tight-junction integrity disrupted by alcohol exposure.

**Figure 2:**
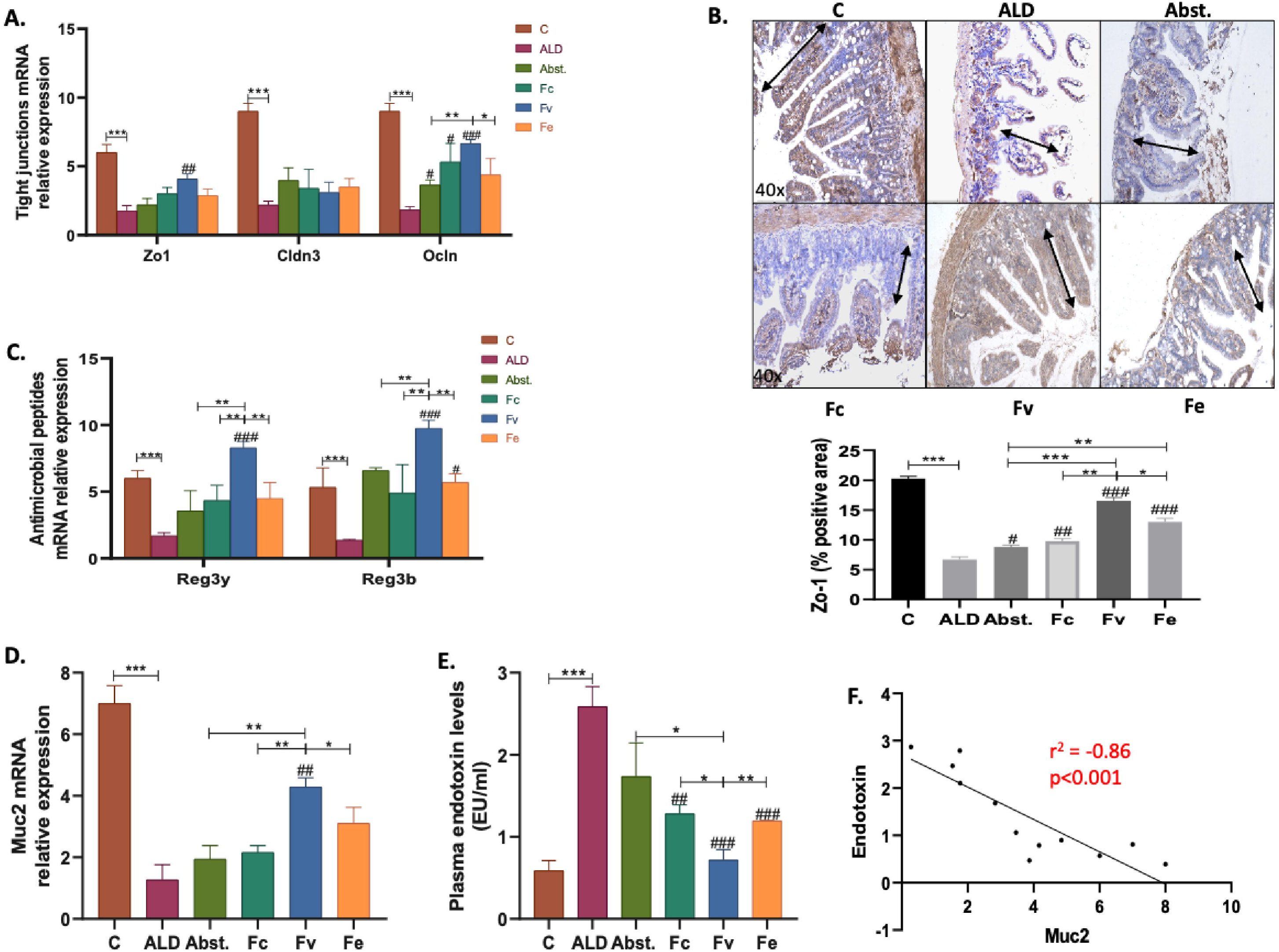
Protein-conditioned FMT restores intestinal barrier integrity. **(A)** mRNA expression of the ileal tight junction proteins Zo-1 (p=0.04), Cldn3(p=0.01), and Ocln (p<0.001) was significantly reduced after Veg-FMT compared to ALD. **(B)** Statistical analysis revealed that the protein expression of the tight junction protein Zo-1 was markedly lower after Veg-FMT than after egg-FMT (1.27FC, p=0.004). (C) mRNA expression of the antimicrobial peptides Reg3ß (p=0.009), Reg3Γ (p=0.006) and **(D)** Muc2 (p=0.05) also decreased after veg-protein conditioned FMT. (**E)** Plasma endotoxin levels (p=0.02) were also significantly lower after veg-protein FMT. **(F)** Serum endotoxin and Muc2 gene expression were significantly negatively correlated (r^2^=-0.86, p<0.001; Spearman’s correlation). The data are presented as the means ± SEMs (standard error of the mean). *p < 0.05, **p < 0.01, ***p < 0.001 represent intergroup statistics; #p < 0.05, ##p < 0.01, ###p < 0.001 with respect to the ALD group (A-E, one-way ANOVA followed by Tukey’s multiple comparison test).

In addition to structural barrier proteins, alcohol exposure also induced intestinal inflammatory responses. Veg-FMT significantly reduced expression of the pro-inflammatory cytokines Il6 (4.2FC decrease, p = 0.002), Tnfα (5.6FC decrease, p < 0.001), and Tlr4 (2FC decrease, p = 0.04) compared with ALD mice, while the anti-inflammatory cytokine Il10 increased 2.8FC (p < 0.001) (Additional file 1, S.Fig. 2B). These findings indicated that Veg-FMT attenuates alcohol-induced intestinal inflammation.

Intestinal antimicrobial defense mechanisms were also affected. Expression of the antimicrobial peptides Reg3β (1.98FC increase, p = 0.01) and Reg3γ (2.04FC increase, p = 0.02) was significantly elevated following Veg-FMT compared with ALD mice (Fig. 2C). Alcohol consumption degrades Muc2 in the gut mucus lining, contributing to a leaky gut and elevated endotoxemia (18). Veg-FMT also significantly increased expression of the mucin gene Muc2 expression (1.4FC, p=0.05; Fig. 2D), with a concomitant reduction in plasma endotoxin levels (1.7FC, p=0.02; Fig. 3E). A significant negative correlation was observed between Muc2 and plasma endotoxin levels (Spearman’s ρ= -0.86, p<0.001; Fig. 2F). These findings indicated that Veg-FMT enhances abstinence-associated restoration of intestinal barrier integrity, leading to reduced gut inflammation and endotoxemia.

**Figure 3:**
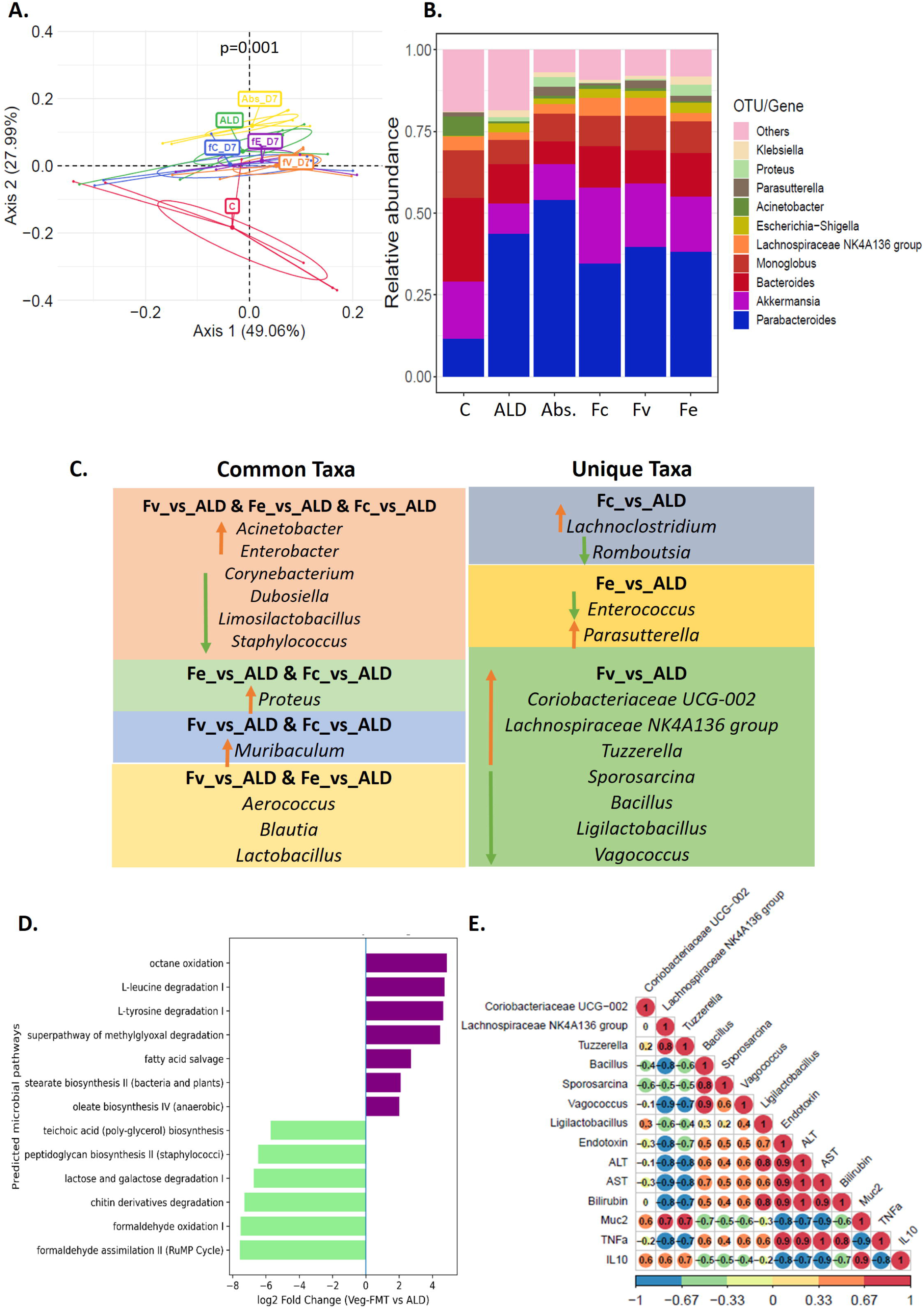
Favorable gut microbial variations were observed after protein-FMT. **(A)** PCoA plot showing significant (p=0.001, PERMANOVA) differences in microbial composition post-FMT among the different treatment groups. **(B)** Comparison of the relative abundances of bacterial genera revealed increases in *Akkermansia, Lachnopiraceae* NK4A136 and *Parasutterella abundance* post-FMT. **(C)** Tables showing differentially altered taxa present commonly and uniquely in the FMT treatment groups. **(D)** Bidirectional bar plot showing selected functional pathways predicted by PICRUSt in veg-FMT **(E)** Correlations between serum injury biomarkers and differential bacterial taxa identified by Veg-FMT. Pearson’s correlation values are displayed in circles. Red represents a positive correlation, whereas blue represents a negative correlation. Correlation according to the scale given at the bottom of the plot.

Given the marked restoration of intestinal barrier integrity following Veg-FMT, we next investigated whether these effects were associated with alterations in gut microbial composition.

### Veg-FMT reshapes gut microbial composition associated with improved metabolic function

Given the significant restoration of intestinal barrier integrity following Veg-FMT, we next investigated whether these improvements were associated with changes in gut microbial composition. To address this, 16S rRNA gene sequencing was performed on fecal samples from six biological animals per experimental group after FMT treatment.

Analysis of alpha diversity using the Shannon index showed no significant differences in overall microbial diversity among the experimental groups (Additional file 1, S.Fig. 3A). However, principal coordinate analysis (PCoA) based on Bray-Curtis distances revealed clear separation between treatment groups, indicating significant differences in microbial community composition (PERMANOVA p = 0.001). The first two principal coordinates explained approximately 77% of the total variance in microbial composition (Fig. 3A).

Taxonomic profiling revealed that Veg-FMT selectively enriched several bacterial taxa associated with beneficial metabolic functions. In particular, the relative abundance of *Akkermansia, Lachnospiraceae* NK4A136 and *Parasutterella* increased following Veg-FMT treatment compared with ALD mice (Fig. 3B). Differential abundance analysis (FC>2, p<0.05) further identified several taxa significantly altered after Veg-FMT (Additional file 1, Fig. S. 3B, C, D, E and Additional file 2, S-table 2A). The abundances of *Coriobacteriaceae UCG-002* (4.1FC, p=0.007)*, Lachnospiraceae* NK4A136 (2.25FC, p=0.02) (SCFA producers that maintain gut barrier function (19)) and *Tuzzerella* (2.45FC, p=0.006; role in lipid metabolism) exclusively increased post-Veg-FMT (Fig. 3C). The opportunistic taxa *Sporosarcina* and *Vagococcus* were exclusively decreased (1.12E-01 FC and 1.11E-02 FC, p<0.001, respectively) suggesting suppression of potentially pathogenic bacterial population. Analysis of taxa shared across treatment groups revealed enrichment of beneficial genera including *Aerococcus, Blautia*, and *Lactobacillus* in protein-modulated FMT recipients.

To further investigate the relationship between microbial changes and host physiology, correlation analysis (r^2^ > 0.7, p<0.05) was performed between differentially abundant bacterial taxa and liver injury parameters (Fig. 3E). Several bacterial taxa enriched after Veg-FMT displayed strong negative correlations with liver injury markers, including serum ALT, AST, bilirubin, and TNFα levels. Notably, *Lachnospiraceae* NK4A136 and *Tuzzerella* showed significant negative correlations with these markers while positively correlating with the intestinal mucus marker Muc2. Together, these findings indicated that Veg-FMT promotes enrichment of beneficial microbial taxa associated with improved intestinal barrier integrity and reduced liver injury. To further explore the potential functional implications of the observed microbial compositional changes, PICRUSt2-based functional inference was performed (Fig. 3D; Additional file 1, Fig. S. F and Additional file 2, S-table 2B). Compared with ALD, Veg-FMT showed significant enrichment of pathways related to amino acid degradation and fatty acid metabolism, including L-leucine degradation (log2FC = 4.74, p = 0.0017), L-tyrosine degradation (log2FC = 4.64, p = 0.0023), octane oxidation (log2FC = 4.86, p = 0.0013), and fatty acid salvage (log2FC = 2.71, p = 0.018). Additional enrichment of lipid biosynthesis pathways such as stearate biosynthesis II (log2FC = 2.09, p = 0.018) and oleate biosynthesis IV (log2FC = 2.01, p = 0.020) further indicated increased microbial fatty-acid metabolic potential. In contrast, several pathways associated with bacterial structural metabolism and carbohydrate degradation, including peptidoglycan biosynthesis II (log2FC = −6.46, p = 0.00023), teichoic acid biosynthesis (log2FC = −5.72, p = 0.0024), and lactose and galactose degradation I (log2FC = −6.74, p = 0.00064), were reduced in Veg-FMT compared with ALD. Collectively, these findings suggest that Veg-FMT is associated with increased predicted microbial metabolic potential, particularly in pathways related to amino acid degradation and fatty acid metabolism.

### Veg-FMT promotes hepatic metabolic reprogramming through PPARα signaling and fatty acid β-oxidation

To determine whether microbiota remodeling induced by Veg-FMT influences hepatic metabolic pathways, we performed quantitative proteomic analysis of liver tissues from control, ALD, and FMT-treated mice.

A total of 3,841 proteins were identified, and unsupervised hierarchical clustering revealed clear segregation between treatment groups, indicating substantial remodeling of hepatic protein expression following FMT (Fig. 4A). Pathway enrichment analysis of the differentially expressed proteins (DEPs; FC > 2, p < 0.05) identified several KEGG pathways associated with metabolic regulation and cellular stress responses (Additional file 1, S. Fig. 4A-E and Additional file 2, S-table 3). The apoptosis pathway, which was elevated in ALD mice, was reversed by all FMT treatments (Additional file 1, S. Fig. 4C-4E). Notably, Veg-FMT uniquely and significantly reduced unsaturated fatty acid biosynthesis (p = 0.04) and showed a marginal reduction in the reactive oxygen species (ROS) pathway (p = 0.08; Additional file 1, S. Fig. 4E).

**Figure 4:**
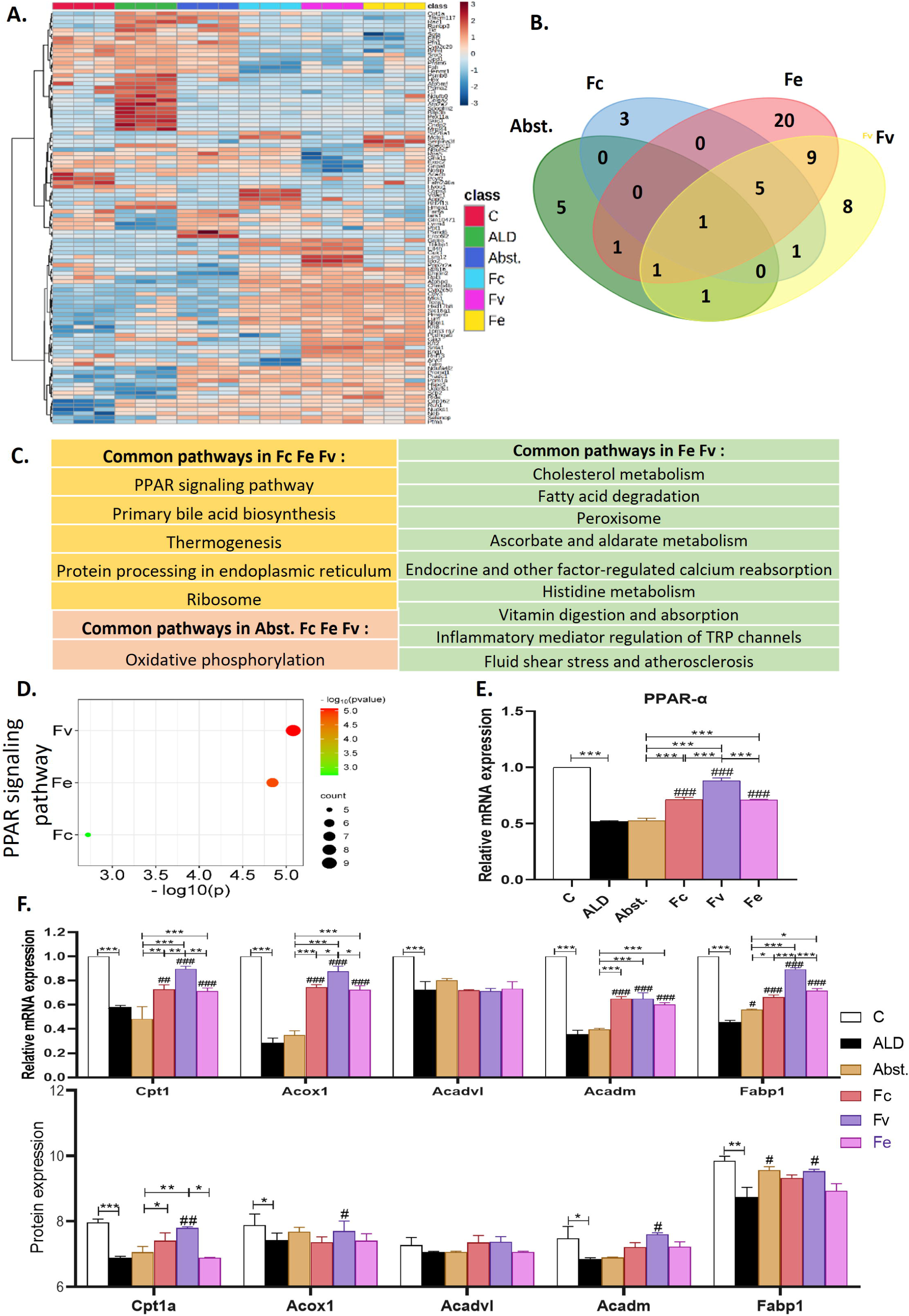
Fecal microbiota transplantation alters the hepatic proteome. **(A)** Heatmap showing differentially expressed proteins whose expression was upregulated or downregulated in different groups. **(B)** Venn diagram showing the number of common and unique enriched pathways among abst, Std-FMT, Veg-FMT and egg-FMT, with the **(C)** table showing pathway names. **(D)** The PPAR signaling pathway was significantly enriched (p=8.3x10-06) in the Veg-FMT group. **(E)** Hepatic PPARα mRNA expression was greater in the Veg-FMT group (p<0.001) than in the Egg-FMT group. **(F)** mRNA expression (top panel; Cpt1 (p=0.003), Fabp1 (p=0.005) and peroxisomal Acox1 (p=0.02)) and protein expression (bottom panel; Cpt1a (p=0.007), Acox1 (p= 0.05), Acadm (p= 0.04) and Fabp1 (p= 0.02)) of PPARα target genes involved in fatty acid beta-oxidation increased after Veg-FMT. The data are presented as the means ± SEMs (standard error of the mean). *p < 0.05, **p < 0.01, ***p < 0.001 represent intergroup statistics; #p < 0.05, ##p < 0.01, ###p < 0.001 represent the ALD group (one-way ANOVA followed by Tukey’s multiple comparison test).

Comparison of common and unique pathways revealed distinct metabolic effects of dietary protein-modulated FMT (Fig. 4B). Stopping alcohol increased oxidative phosphorylation, whereas all FMT treatments significantly enriched pathways related to PPAR signaling, primary bile acid biosynthesis, and endoplasmic reticulum protein processing. In addition, protein-modulated FMT specifically enhanced pathways associated with cholesterol metabolism, fatty acid degradation, and histidine metabolism (Fig. 4C, Additional file 2, S-table 4).

Among the enriched pathways, PPAR signaling emerged as a prominent metabolic pathway associated with FMT-mediated recovery. Given the well-established role of peroxisome proliferator–activated receptors (PPARs) in regulating hepatic lipid metabolism and steatosis (20,21), we further examined proteins associated with this pathway. Enrichment analysis identified 9 proteins linked to PPAR signaling in Veg-FMT (p = 8.3 × 10 ), 7 proteins in Egg-FMT (p = 1.44 × 10 ), and 5 proteins in Std-FMT (p = 0.001; Fig. 4D). Among these, PPARα expression was significantly increased following Veg-FMT (1.7FC vs ALD, p < 0.001; 1.2FC vs Egg-FMT, p < 0.001) (Fig. 4E), whereas PPARγ expression remained unchanged across treatment groups (Additional file 1, S.Fig. 4F).

To determine whether PPARα activation translated into functional metabolic changes, we examined downstream genes involved in fatty acid β-oxidation. Both Veg-FMT and Egg-FMT increased β-oxidation–related gene expression; however, Veg-FMT produced significantly stronger effects. Compared with Egg-FMT, Veg-FMT significantly increased expression of mitochondrial Cpt1 (1.25FC, p = 0.003), Fabp1 (1.25FC, p = 0.005), and peroxisomal Acox1 (1.2FC, p = 0.02; Fig. 4F). Consistent with these findings, protein expression of Cpt1a (1.2FC, p = 0.007), Acox1 (1.1FC, p = 0.05), Acadm (1.1FC, p = 0.04), and Fabp1 (1.1FC, p = 0.02) was significantly increased only in Veg-FMT–treated mice; Fig. 4F). Thus, Egg-FMT also engaged PPARα-associated fatty acid metabolic responses, but the magnitude of these responses was consistently lower than that observed with Veg-FMT. Together, these findings indicated that vegetable protein–modulated FMT more effectively activates hepatic PPARα signaling and downstream fatty acid β-oxidation pathways, indicating enhanced hepatic lipid metabolism following Veg-FMT treatment.

### Veg-FMT mediated metabolic effects are dependent on PPARα signaling

To determine whether the metabolic benefits of Veg-FMT were mediated through PPARα signaling, we pharmacologically inhibited PPARα using the antagonist GW6471 in Veg-FMT–treated ALD mice (Fig. 5A). PPARα inhibition significantly reduced hepatic PPARα mRNA expression (4.5FC decrease, p < 0.001; Fig. 5B) and protein levels (2.3FC decrease, p = 0.01; Fig. 5C) compared with Veg-FMT alone. Consistent with suppression of this pathway, inhibition of PPARα resulted in a significant increase in the liver-to-body weight ratio (3.3FC, p = 0.04; Additional file 1, S.Fig. 5A).

**Figure 5:**
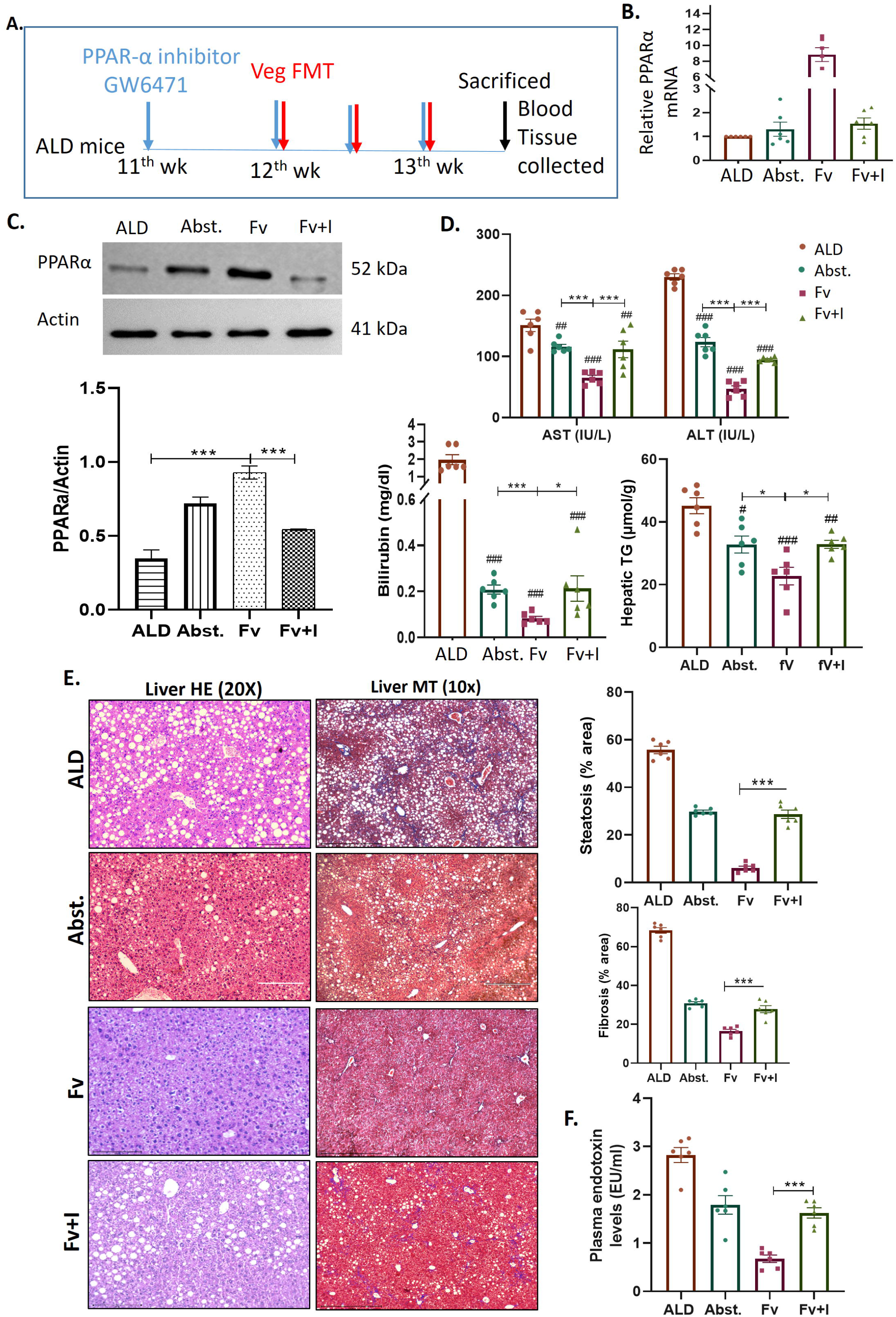
Figure 5**: Veg-FMT alleviates hepatic injury through PPAR**α **activation. (A)** Study design showing PPARα inhibitor administration in ALD mice with Veg-FMT. At 7 days after FMT, the tissues and blood were collected for further analyses. **(B)** mRNA (p<0.001) and **(C)** protein expression of PPARα decreased significantly (p=0.01). **(D)** Serum AST(p=0.003), ALT(p=0.002), bilirubin(p=0.04) and hepatic triglyceride levels (p=0.03) significantly increased after PPARα inhibition. **(E)** Liver histology images showing increased steatosis (p<0.001) by H&E and fibrosis(p<0.001) by MT staining after inhibition copared to veg-FMT. **(F)** Plasma endotoxin levels (p=0.001) were also increased after inhibition. The data are presented as the means ± SEMs (standard error of the mean). *p < 0.05, **p < 0.01, ***p < 0.001 represent intergroup statistics; #p < 0.05, ##p < 0.01, ###p < 0.001 with respect to the ALD group (one-way ANOVA followed by Tukey’s multiple comparison test).

Systemic markers of liver injury were also markedly elevated following PPARα inhibition, including serum AST (1.7FC, p = 0.003), ALT (2FC, p = 0.002), and bilirubin (2.5FC, p = 0.04), accompanied by increased hepatic triglyceride accumulation (1.45FC, p = 0.03; Fig. 5D). Histological analysis further revealed persistence of hepatic inflammation, steatosis, and fibrosis in the inhibitor-treated group (Additional file 1, S.Fig. 5C; Fig. 5E), indicating loss of the hepatoprotective effects observed with Veg-FMT alone.

PPARα inhibition also impaired intestinal barrier integrity, as evidenced by villus blunting, reduced ZO-1 protein expression (1.97FC decrease, p < 0.001; Additional file 1, S.Fig. 5D), and significantly elevated plasma endotoxin levels (2.3FC increase, p < 0.001; Fig. 5F). Consistently, expression of canonical PPARα target genes involved in fatty acid β-oxidation, including Cpt1, Acox1, and Fabp1, was markedly suppressed (5FC, 4.3FC, and 4.7FC decreases, respectively; p < 0.001 for all; Additional file 1, S.Fig. 5E).

Interestingly, inhibition of PPARα did not significantly alter overall gut microbiota composition between groups (PERMANOVA p = 0.644; Additional file 1, S.Fig. 6), suggesting that the loss of protection was primarily due to disruption of host metabolic signaling rather than changes in microbial community structure.

**Figure 6:**
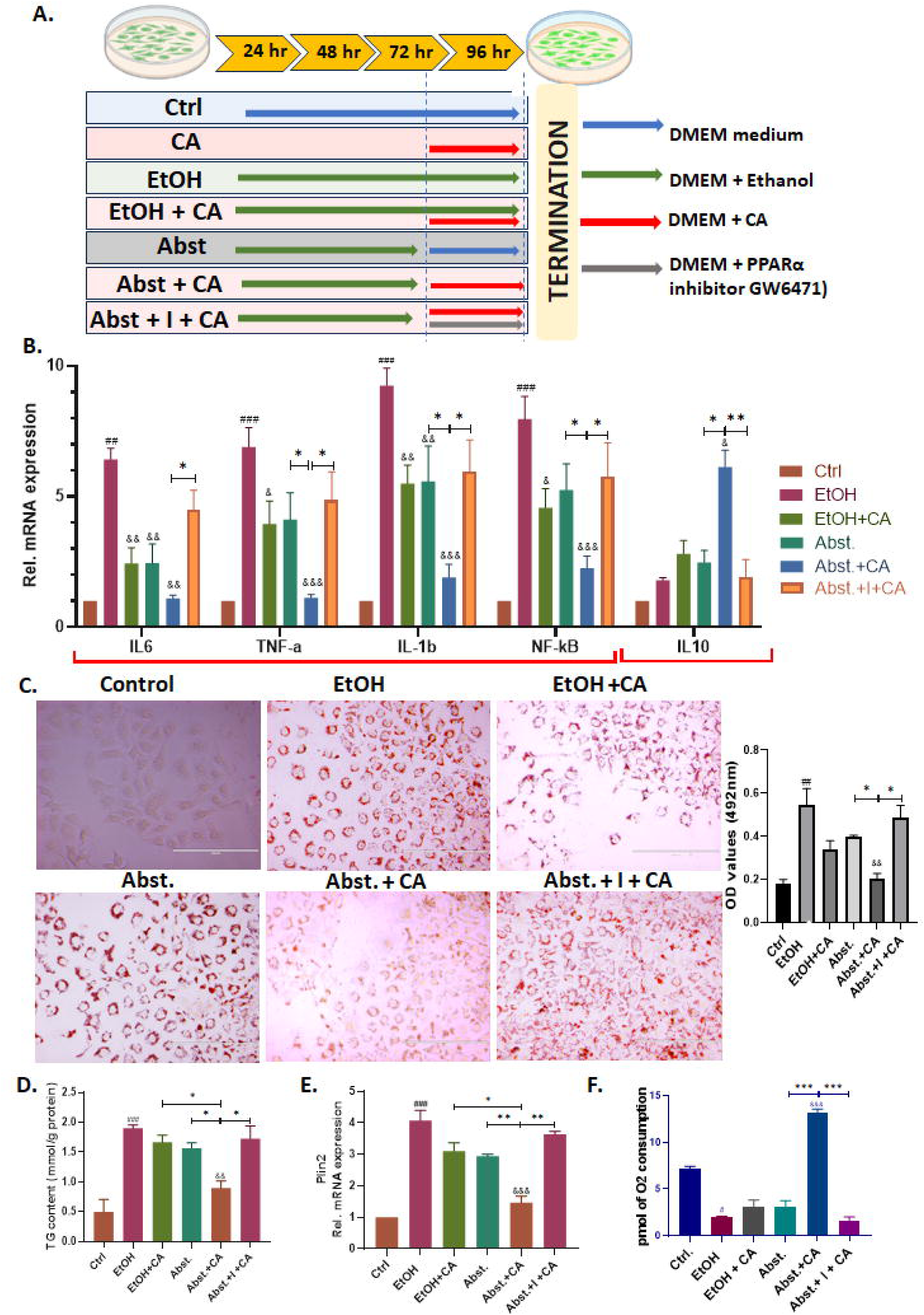
Caproic acid supplementation mitigates ethanol-induced steatosis by enhancing fatty acid. β**-oxidation in Huh7 cells. (A)** Study design: Huh7 cells were treated with ethanol or caproic acid alone or in combination. A PPARα inhibitor was also given with caproic acid. Injury was assessed after 24 hours. **(B)** mRNA expressions of pro-inflammatory reduced (IL6, p=0.002; TNFα, p=0.006; IL1β, p<0.001; Nfkb, p<0.001) and anti-inflammatory markers (IL10, p=0.005) increased after Abst+CA supplementation and these changes reversed in presence of PPARα inhibitor. **(C)** Cytopathology showing higher lipid accumulation in ethanol treated cells using oil-red-o staining and significantly reduced after CA supplementation (p=0.02). CA supplementation effects were reduced in the presence of PPARα inhibitor (p=0.01). **(D)** TG levels were significantly decreased after CA supplementation (p=0.05) and effects were reversed after PPARα inhibition. (E) mRNA expression of lipid droplet marker (Plin2) increased in ethanol treatment and reduced CA supplementation (p=0.002), but reversed in presence of PPARα inhibitor. **(F)** Mitofuel flexibility assay showed increased β-oxidation dependency after CA supplementation (p<0.001), but it was reduced in the presence of the inhibitor (p<0.001). The data are presented as the means ± SEMs (standard error of the mean). *p < 0.05, **p < 0.01, ***p < 0.001 represent intergroup statistics; #p < 0.05, ##p < 0.01, ###p < 0.001 with respect to the ALD and control groups. &p < 0.05, &&p < 0.01, &&&p < 0.001 with respect to ALD (one-way ANOVA followed by Tukey’s multiple comparison test).

Collectively, these findings indicated that activation of hepatic PPARα signaling is essential for the protective metabolic effects of Veg-FMT, linking microbiota-derived metabolic signals to host lipid oxidation and liver recovery.

### Metabolomic profiling identifies caproic acid enrichment following Veg-FMT

PPARα has multiple metabolic ligands (22). Untargeted metabolomic profiling was performed on stool and plasma samples to identify microbiota associated metabolites potentially involved in PPARα signaling. A total of 665 metabolites in stool and 522 in plasma were detected. Principal coordinate analysis indicated clear separation between experimental groups, with the first two components explaining ∼60% of the variance in stool metabolites and ∼31% in plasma metabolites (Additional file 1, S. Fig. 7A-B). Differential metabolite analysis revealed multiple metabolites significantly altered in Abst- and FMT-treated animals compared with ALD mice (FC>1.5, p<0.05) (Additional file 1, S. Fig. 7C-D; Additional file 2, S-table 5 for stool and S-table 6 for plasma). Unique significantly upregulated metabolites for both stool and plasma were identified (Additional file 1, S. Fig. 7E-F; Additional file 2, S-table 7 for stool and 8 for plasma). Comparative analysis of protein-modulated FMT groups showed 72 metabolites commonly increased in stool and 7 in plasma, whereas Veg-FMT uniquely increased 61 stool and 23 plasma metabolites. Among these metabolites, two molecules were significantly enriched in both stool and plasma following Veg-FMT: N-butanoyl-homoserine lactone and caproic acid (Additional file 1, S. Fig. 7G and Additional file 2, S-table 9). N-butanoyl-homoserine lactone is a bacterial quorum-sensing molecule involved in microbial communication and biofilm regulation (23). In contrast, caproic acid (hexanoic acid) is a microbial fatty acid metabolite associated with anti-inflammatory effects and host energy metabolism (24). Caproic acid levels were significantly increased in Veg-FMT recipients in both stool (2.5FC, p=0.01) and plasma (3.4FC, p = 0.01), and were higher compared with Egg-FMT animals (Additional file 1, S. Fig. 7H-I). Notably, Egg-FMT also showed an increase in caproic acid, although at lower levels than Veg-FMT, consistent with its intermediate hepatic phenotype.

**Figure 7:**
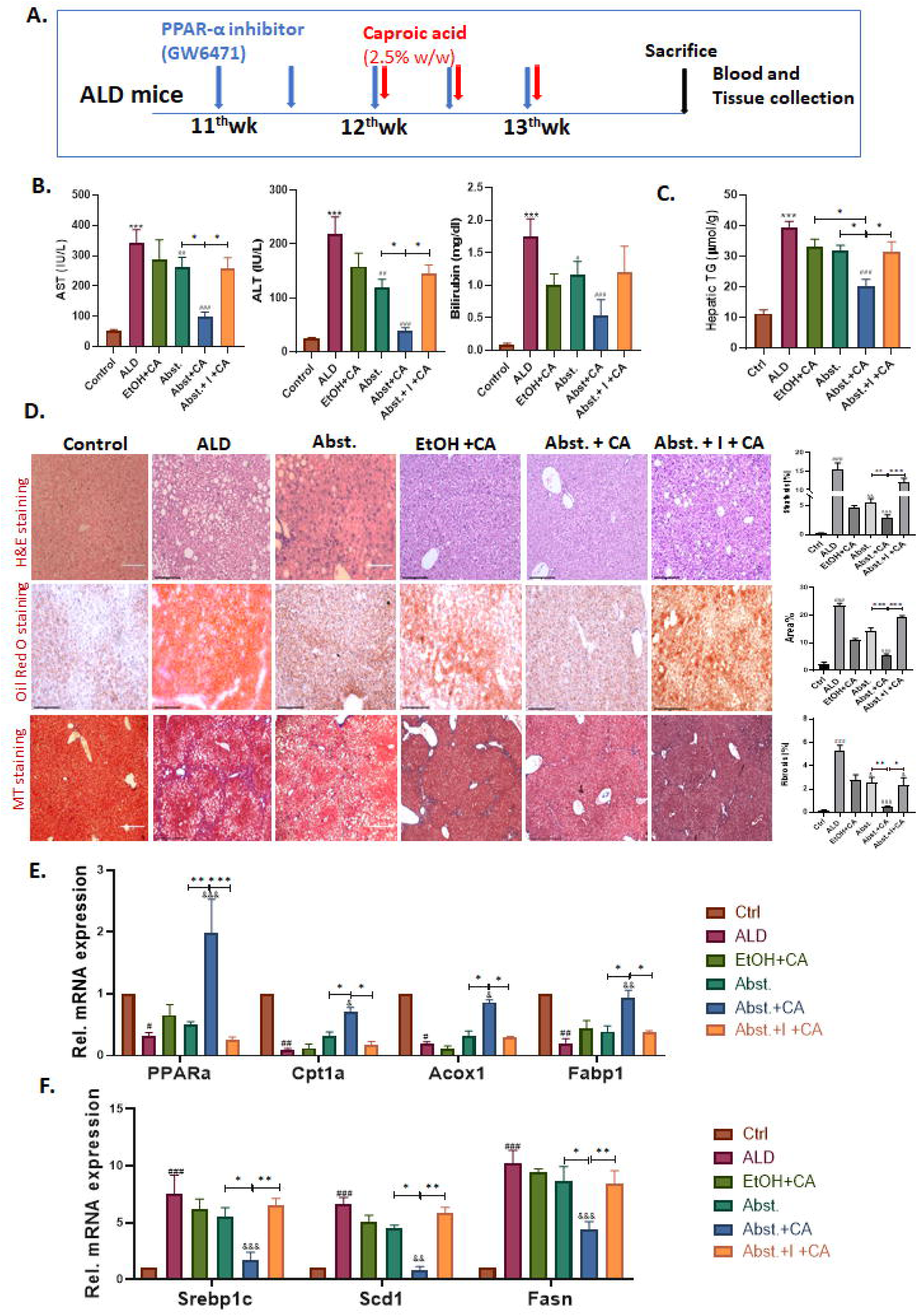
**Caproic acid supplementation alleviated ethanol-induced liver injury through PPAR**α **upregulation in ALD mice. (A)** Study design-ALD mice treated with caproic acid with or without a PPARα inhibitor. Injury was assessed after 1 week. **(B)** Serum AST (p=0.02), ALT (p<0.001) and bilirubin (p=0.006) levels were significantly reduced after CA supplementation. **(C)** Hepatic TGs were significantly reduced after CA supplementation (p=0.02) and these changes were reversed in the presence of PPARα inhibitor **(D)** Liver histology showing reduced steatosis (examined by H&E and Oil Red O, p<0.001 in both) and fibrosis (p=0.007) by MT staining after CA supplementation. **(E)** mRNA expression of PPARα target genes involved in fatty acid β-oxidation increased (PPARα, p<0.001; Cpt1, p=0.01; Acox1, p=0.008; Fabp1, p=0.002) and **(F)** lipogenesis-related mRNA expression decreased (Srebp1c, Scd1 and Fasn, p<0.001 in all) after CA supplementation. These beneficial effects of CA were attenuated in the presence of a PPARα inhibitor. The data are presented as the means ± SEMs (standard error of the mean). *p < 0.05, **p < 0.01, ***p < 0.001 represent intergroup statistics; #p < 0.05, ##p < 0.01, ###p < 0.001 with respect to the ALD and control groups. &p < 0.05, &&p < 0.01, &&&p < 0.001 with respect to ALD (one-way ANOVA followed by Tukey’s multiple comparison test).

These findings identified caproic acid as a candidate microbiota-derived metabolite potentially linking Veg-FMT induced microbiota remodeling with hepatic metabolic responses.. To further examine the relationship between microbiota-associated metabolites and host responses, we performed an integrated Spearman correlation analysis using matched microbiome, metabolomic, and phenotypic data (Additional file 1, Fig. S8; S. Table 10). *Lachnospiraceae* NK4A136 and *Coriobacteriaceae* UCG-002 were positively associated with caproic acid abundance (ρ=0.713 and 0.678, respectively; FDR q<0.05). Caproic acid was negatively associated with hepatic triglycerides (ρ= -0.797, q=0.006), AST (ρ= -0.779, q=0.008), bilirubin (ρ =-0.764, q=0.009), IL-6 (ρ= -0.763, q=0.009), and ALT (ρ= -0.615, q=0.043). These findings provide associative support for a microbiota-caproic acid-host metabolic axis and supported the selection of caproic acid for subsequent functional validation, while not establishing causality.

### Caproic acid mitigates ethanol-induced steatosis by enhancing fatty acid β-oxidation in hepatocytes

Because fatty acids can function as PPARα ligands (25), we next examined whether caproic acid could modulate PPARα signaling in hepatocytes. Ethanol-exposed Huh7 cells were treated with caproic acid (CA) in the presence or absence of the PPARα inhibitor GW6471 (Fig. 6A). Based on dose-response experiments, 100 mM ethanol and 900 µM caproic acid were selected for subsequent studies (Additional file 1, S.Fig. 9A).

CA supplementation significantly reduced ethanol-induced inflammatory responses, including IL-6 (5FC, p = 0.002) and TNFα (5.5FC, p = 0.006), while increasing the anti-inflammatory cytokine IL-10 (Fig. 6B). These effects were reversed in the presence of GW6471, indicating that the anti-inflammatory effects of CA are mediated through PPARα signaling.

Consistent with this observation, CA markedly suppressed lipogenic gene expression, including Srebp1c (4FC, p < 0.001) andFasn (4.5FC, p < 0.001), while significantly increasing β-oxidation genes, including PPARα (8FC, p < 0.001), Cpt1a (7.6FC, p = 0.001), and Fabp1 (5.7FC, p = 0.002) (Additional file 1, S. Fig. 9B–C). Oil Red O staining confirmed a marked reduction in lipid accumulation following CA treatment (2FC decrease, p = 0.02), which was reversed by PPARα inhibition (Fig. 6C).

Consistently, intracellular triglyceride levels were significantly reduced after CA supplementation (1.7FC decrease, p = 0.05; Fig. 6D), accompanied by reduced expression of the lipid droplet marker PLIN2 (2.07FC decrease, p = 0.002; Fig. 6E). In contrast, genes involved in lipid export (ApoB and MTTP; Additional file 1, S. Fig. 9D) showed only modest and non-significant changes, suggesting that CA-mediated steatosis reduction primarily results from enhanced fatty acid oxidation rather than increased lipid export.

To functionally assess alterations in cellular fatty acid utilization, a Mito Fuel Flex assay was performed. CA significantly increased oxygen consumption associated with fatty acid oxidation (4.2FC, p < 0.001; Fig. 6F), and this effect was completely abolished by GW6471. In comparison, the known PPARα ligand butyric acid (25) induced only a modest increase in fatty acid oxidation (Additional file 1, S. Fig. 9E). Because the Mito Fuel Flex assay assesses cellular substrate dependency and utilization rather than directly quantifying intracellular β-oxidation flux, these findings were interpreted as functional evidence of altered fatty acid utilization rather than a direct measurement of metabolic flux. To further analyse the binding kinetics between PPARα and Caproic acid, we performed biolayer interferometry using recombinant PPARα ligand-binding domain (PPARα-LBD). Caproic acid showed direct concentration-dependent association curves for the increasing concentrations of CA (31.25–500 µM). Each reaction displayed clean association and dissociation curves in 1X PBS as reaction buffer mimicking cellular milieu. Affinity constant of 46µM±7.6E-21M, strengthens our validation for direct interaction between PPARα and Caproic acid (Additional file 1, Fig. S10A-B). These findings provide complementary biophysical evidence for direct interaction between caproic acid and the PPARα ligand-binding domain. Together, these findings support a role for PPARα signaling in the caproic acid-associated reduction of hepatocellular lipid accumulation and altered fatty acid utilization.

### In vivo confirmation of caproic acid-mediated hepatoprotection

To validate these findings in vivo, ALD model mice were supplemented with caproic acid with or without the PPARα inhibitor (Fig. 7A). CA supplementation significantly reduced the liver-to-body weight ratio compared with ethanol-treated mice (1.5 FC, p<0.001; Additional file 1, S. Fig. 11A), CA supplementation also significantly reduced serum markers of liver injury, including AST (3.4FC, p = 0.02), ALT (5.5FC, p < 0.001), and bilirubin (10FC, p = 0.006), whereas these protective effects were completely abolished in the presence of the inhibitor (Fig. 7B).

Histological and biochemical analyses further indicated significant reductions in hepatic lipid accumulation following CA supplementation. Hepatic triglyceride levels decreased significantly (1.6FC, p = 0.02; Fig. 7C), accompanied by marked reductions in H&E-positive steatotic area (5FC, p < 0.001) and Oil Red O-positive lipid accumulation (4.3FC, p < 0.001). CA supplementation also significantly reduced hepatic fibrosis (5FC, p = 0.007; Fig. 7D), reduced hepatic expression of pro-inflammatory markers (IL6, TNFa, IL1b, NFkB and increased the expression of the anti-inflammatory marker (IL10; Additional file 1, S. Fig. 11B). These protective effects were not observed in mice treated with the PPARα inhibitor.

At the molecular level, CA significantly suppressed lipogenic genes, including Srebp1c (4.4FC), Scd1 (7.3FC), and Fasn (2.3FC; p < 0.001 for all), while simultaneously enhancing expression of PPARα and downstream β-oxidation genes, including Cpt1 (7FC, p = 0.01), Acox1 (4.7FC, p = 0.008), and Fabp1 (4.8FC, p = 0.002) (Fig. 7E-F). These transcriptional changes were attenuated upon PPARα inhibition, while the overall hepatoprotective effects of CA were also diminished, supporting a functional role for PPARα signaling in the response to CA. Collectively, these results suggested that caproic acid supplementation mitigates alcohol-induced liver injury by activating PPARα-mediated fatty acid β-oxidation pathways in vivo.

## DISCUSSION

The current study identifies veg-protein enriched donor diet followed by FMT (veg-FMT) as a therapeutic option for ameliorating alcohol-associated liver disease (ALD). We identified caproic acid, a six-carbon fatty acid produced by gut bacteria (26), as a candidate microbial mediator associated with PPARα-dependent regulation of hepatic fatty acid metabolism.. A key finding is the significant role of donor preparation, particularly protein supplementation and the kind of protein in the diet, which enhances the efficacy of FMT. By modulating the donor gut microbiota through plant-based proteins, Veg-FMT accelerated and amplified abstinence related recovery, leading to reduced hepatic inflammation and fibrosis-associated markers and staining..

Veg-FMT also significantly enhanced intestinal barrier integrity with a concomitant reduction in plasma endotoxin levels, highlighting its dual action in mitigating gut epithelium damage and limiting systemic inflammation. These findings are consistent with previous studies showing that microbial interventions restore gut permeability and reduce endotoxemia in ALD models (7,27,28).

Additionally, the upregulation of antimicrobial peptides (Reg3β and Reg3γ) highlights the role of Veg-FMT in strengthening mucosal immunity. These peptides likely contribute to maintaining a healthier gut microbiota composition, which is essential for reducing alcohol-induced endotoxemia (29). These findings suggest that the benefits of Veg-FMT rely on preserving intestinal barrier integrity and mucosal defenses, which is consistent with our earlier work showing that plant protein supplementation improved hepatic stress and barrier function in ALD (30).

Donor management plays a pivotal role in the success of FMT (31). Given that diet directly influences the composition of the gut microbiota (12), numerous studies have highlighted the impact of pre-educating donor microbiota with different dietary regimens on the efficacy of FMT (32–34). Kedia et al. reported that FMT plus an anti-inflammatory diet significantly improved UC outcomes, with deep remission maintained for up to a year through ongoing dietary interventions (35). Similarly, Rinott et al. reported that diet-modulated autologous FMT, especially from green-Mediterranean diet donors, reduced weight regain and improved glycemic control, highlighting the lasting metabolic benefits of diet-driven microbiome modulation (36). In addition, Yang et al. suggested that FMT from methionine-restricted diet donors remodeled the gut microbiota in obese mice, increasing SCFA-producing bacteria, reducing proinflammatory taxa, and improving lipid metabolism and fat browning (33). Together, these findings highlight the importance of donor preparation, especially through tailored diets, to increase the therapeutic potential of FMT.

In addition to broader dietary regimens, specific dietary macronutrients, such as proteins, further influence the composition of the gut microbiota (11). While dietary fiber and prebiotic oligosaccharides are established substrates for gut fermentation, emerging evidence shows that small amounts of nondigestible dietary proteins and peptides also reach the colon for microbial metabolism. The differing digestibility of plant- and animal-based proteins shapes their distinct effects on the gut microbiota, with plant proteins, due to lower digestibility and complexity, being more likely to reach the colon and drive unique microbial profiles (37). Egg albumin was selected as the animal-derived protein source because of its well-characterized composition, high biological value, and prior use in our ALD dietary intervention studies (30). However, egg albumin represents only one animal protein source, and other proteins such as casein or whey may produce distinct microbial and metabolic responses. Therefore, our findings should not be generalized to animal proteins as a whole.

In this context, Veg-FMT, which involves plant protein-modulated microbiota, results in a compositional shift in the gut microbiota, characterized by an increased abundance of beneficial taxa such as *Akkermansia, Lachnospiraceae* NK4A136, and *Parasutterella*. *Lachnospiraceae* NK4A136 and *Tuzzerella* are known SCFA producers and mediate anti-inflammatory pathways (19). *Akkermansia* has been shown to maintain the mucus layer in the gut (38). These taxa have been associated with improved intestinal barrier function and reduced systemic inflammation. PICRUSt2-based functional inference suggested enrichment of microbial pathways related to amino acid degradation and fatty acid metabolism following Veg-FMT. These predictions were accompanied by measurable changes in microbial associated metabolites, including increased caproic acid, providing observational support for altered microbial metabolic potential. (39,40).

Changes in the gut microbiota can influence liver function. Our proteomics and pathway analyses revealed that Veg-FMT significantly activated PPARα signaling, a key regulator of impaired fatty acid oxidation in ALD (41). The upregulation of PPARα and its targets (Cpt1, Acox1, and Fabp1) enhanced mitochondrial and peroxisomal β-oxidation, reducing hepatic lipid accumulation. Pharmacological inhibition of PPARα attenuated the protective metabolic effects of Veg-FMT, supporting a functional role for PPARα signaling in these responses.. These results align with those of previous studies highlighting PPARα as a therapeutic target in ALD (42,43).

Microbial mediators derived from dietary protein fermentation may modulate PPARα signaling. In the colon, microbial fermentation of dietary proteins generates SCFAs and other metabolites (39). Among these metabolites, caproic acid, a six-carbon fatty acid, emerged from our metabolomic analysis as a candidate mediator of PPARα-associated metabolic responses. Fatty acids can interact directly with PPARα and modulate its activity (44). Consistently, previous studies have shown that short- and medium-chain fatty acids contribute to cellular energy metabolism and can influence lipid metabolic pathways (24). In vitro and in vivo supplementation with caproic acid reduced hepatic lipid accumulation and altered fatty acid utilization, while these effects were attenuated by PPARα inhibition. Consistent with the established ability of fatty acids to interact with PPARα, BLI analysis demonstrated direct interaction between caproic acid and the PPARα ligand-binding domain, providing complementary biophysical evidence for PPARα engagement by caproic acid. Together, these findings support a role for caproic acid in PPARα-dependent hepatic metabolic responses. Notably, caproic acid-mediated improvement in hepatic lipid homeostasis was also associated with reduced triglyceride accumulation and downregulation of the lipid droplet-associated protein PLIN2, whereas lipid export genes (ApoB and MTP) were only modestly or non-significantly altered, indicating that steatosis attenuation was driven primarily by reduced lipid storage and enhanced fatty acid β-oxidation rather than increased VLDL export (45,46).

However, the study has several limitations. Findings are based on an animal model and require validation in human studies for FMT application. The long-term safety and broader metabolic impacts of Veg-FMT remain unknown. The modest sample size and inter-individual variability across omics analyses may limit statistical power and increase the risk of false-positive associations; therefore, these findings should be interpreted cautiously. Furthermore, individual microbiota variability could influence treatment reproducibility, highlighting the need for personalized approaches in future research. The one-week post-FMT observation period was designed to capture early therapeutic and mechanistic responses but is insufficient to establish durable fibrosis reversal or long-term structural remodeling; longer-term studies are needed to determine the persistence of these effects. Short-course FMT has shown beneficial outcomes in patients with ALD, including a seven-day FMT course in severe alcoholic hepatitis (8,47) and a single FMT session in alcohol-associated ACLF (48), with a meta-analysis supporting improved short-term survival (49). However, although we observed a rapid reduction in fibrosis after FMT in this animal model, the response is likely to be slower in patients. In clinical studies, liver stiffness changed over one week to one month in alcohol-associated cirrhosis without a change in fibrosis grade (50). Fibrosis improvement in patients may therefore take weeks to months. Given the role of PPARα in ALD, pharmacological activation of this pathway could be explored as a therapeutic strategy. Elafibranor, a dual PPARα/δ agonist, has shown promise in improving liver function and reducing inflammation in ALD (51).

## Conclusion

This study provides new insights into the mechanisms underlying the therapeutic effects of Veg-FMT. We show that preconditioning the donor microbiota with a plant-protein diet is key to enhancing FMT efficacy by promoting SCFA-producing bacteria and elevating caproic acid levels. Caproic acid, in turn, activates PPARα, increasing β-oxidation, suppressing lipogenesis, and improving hepatic injury and metabolic health. To our knowledge, this is the first demonstration of the specific pathways that mediate the benefits of FMT.

## MATERIALS AND METHODS

### Adaptation of mice to high-protein diets

Male C57BL/6 N mice (strain code-632) aged 6-8 weeks and weighing 20-25 g were procured from the Center for Comparative Medicine, ILBS, Delhi. The animals were housed under barrier housing conditions following specific pathogen-free animal husbandry practices. The ambient temperature and humidity were maintained at 21–25°C and 50–70%, respectively. The Institutional Animal Ethics Committee approved the animal protocols, and the experiments were performed following the ARRIVE (Animal Research: Reporting of In Vivo Experiments) guidelines 2.0. The animal experiments were performed in the Experimental Animal Facility of ILBS, Delhi, during the light cycle. Autoclaved corn cob bedding material and a ventilated cage system were used for the housing. A piece of M-block wood (Kansara Scientific, India) was supplied to the mice so that they could gnaw on it to reduce stress.

The animals were acclimatized to a liquid diet for 1 week. Mice were then fed incremental alcohol in Lieber-DeCarli liquid diet (Cat#F1258SP, Bio-Serv, USA) with thioacetamide for 12 weeks to develop the ALD model as described earlier (52). Control mice were pair-fed a non-alcoholic isocaloric Lieber-DeCarli liquid diet (Cat#F1259SP, Bio-Serv, USA; calories equated with maltodextrin). For FMT the donor mice were randomly divided into three groups: control diet donor, high vegetable protein diet donor (crushed soya chunks; RSI Ltd., India), and egg protein diet donor (albumin egg flakes; Cat#GLR09.020738). All the diets were kept isocaloric (Additional file 1, S.Fig 1A). Respective diets were given for 14 days, and their fecal matter was collected at day 14. FMT recipients were fed standard liquid diet for seven days prior to FMT. The measurement of body weight was conducted every week for each animal, whereas food intake was assessed once every two days.

### Fecal microbiota transplantation

For the FMT study, 6 mice were used per donor group (three donor groups: Cd, Vd and Ed). ALD was established in four groups of animals (n=8 each): Abstinence (Abst), standard FMT (Std-FMT), veg-protein FMT (Veg-FMT), and egg-protein FMT (Egg-FMT). The stool from animal donors fed high-protein or standard diets was collected and immediately processed for FMT. Fresh stool (300 mg) was resuspended in 3 ml of sterile, ice-cold 0.9% saline. Recipient ALD mice received gut washes with 200 µl of PEG-4000 (Cat#81240, Sigma Aldrich, USA). After four hours, 200 μl of the prepared fecal slurry was administered via oral gavage. Three doses of FMT were given every other day. The experiment was terminated 1 week after the last FMT dose was given.

### Tissue collection

The stool was collected pre- and post-FMT. All other tissue samples were collected at the end of the experiments. Blood was obtained via retro-orbital sinuses just before euthanasia (Ketamine, 80-100mg/Kg + Xylazine, 8-10 mg/Kg). Tissue samples were weighed and divided into two groups – one set snap-frozen in liquid nitrogen, and stored at −80°C till further processing. The second group for histopathology was fixed in 10% formalin.

### Biochemical tests

To investigate the effect of FMT, serum was separated from the blood by centrifugation at 3,000xg for 10 minutes. Alanine aminotransferase (ALT), aspartate aminotransferase (AST), bilirubin, and triglycerides were detected by an Olympus AU5400 automatic analyzer (Olympus Optical, Tokyo, Japan). Serum endotoxin levels were measured per manufacturer’s protocol using Pierce Chromogenic Endotoxin Quant Kit (#A39552, Thermofisher Scientific).

### Histopathology and immunohistochemistry

Post-FMT histopathological features of liver and ileum were assessed in the formalin fixed tissue sections (4 μm thick) stained with hematoxylin-eosin (H&E) to capture immune infiltrations, steatosis (Oil red O staining) and fibrosis (Masson’s trichrome staining). Immunohistochemical staining was performed to assess extent of fibrosis using antibodies against α-SMA (#MA5115467, Thermo Fischer, USA) on hepatic tissue. The sections were washed and incubated with an appropriate biotinylated secondary antibody and then with streptavidin-horseradish peroxidase complex (PolyExcel HRP/DAB Detection System Two Step Universal Kit, #PEH002, PathnSitu Biotechnologies, USA). Sections were then counterstained with hematoxylin (#S034, HiMedia). The sections were observed with the help of a light microscope (EVOSTM M5000, Thermofisher Scientific, USA) at a magnification of 10x, 20x and 40x.

Additional details of methods and materials can be found in supplementary section 1 (S1). **Abbreviations:** Acadm: Acyl-Coenzyme A dehydrogenase, medium chain; Acadvl: Acyl-Coenzyme A dehydrogenase, very long chain; Acox1: Acyl-Coenzyme A oxidase 1; Acta2: actin alpha 2, smooth muscle; ALT: alanine aminotransferase; ALD: Alcohol-related liver disease; APOB: Apolipoprotein B; AST: aspartate aminotransferase; Cldn3: claudin3; Cpt1: Carnitine palmitoyltransferase 1; Fabp1: Fatty acid binding protein 1; Fasn: fatty acid synthase; Fc: Std-FMT; Fe:Egg-FMT; FMT: Fecal microbiota transplant; Fv:Veg-FMT; IL: Interleukin; MTTP: MTTP: Microsomal Triglyceride Transfer Protein; Muc2: mucin 2; Ocln: occludin; PCoA: principal coordinate analysis; PLIN2: Perlipin 2; PPARα: peroxisome proliferator-activated receptor alpha; Scd1: stearoyl-coenzyme A desaturase 1; Srebp1c: Sterol Regulatory Element Binding Protein 1c; Tgf-β: tumor growth factor beta; Tlr4: toll-like receptor 4; Tnf-α: tumor necrosis factor alpha. TGs: Triglycerides.

## Availability of data and material

The 16s fastq data have been deposited with links to BioProject accession number PRJDB35434 in the DDBJ BioProject database. The raw metabolomic files can be accessed from NGDC OMIX010622 under Bioproject PRJCA041599. The mass spectrometry proteomics data have been deposited to the ProteomeXchange Consortium via the PRIDE partner repository with the dataset identifier PXD064611. The reviewer can access the dataset by logging into the PRIDE website via the following account details: Username, Password: M9ehP1lOAcK2

## S-12/1/2021-SCHEME (CoE)

## Author Contributions

**Nishu Choudhary#:** Conceptualization; Data curation; Formal analysis, Methodology, Validation; Visualization; Writing – original draft, review & editing; **Ashi Mittal#:** Conceptualization; Data curation; Formal analysis, Methodology, Validation; Visualization; Writing – original draft, review & editing; **Sandeep Kumar:** Methodology, Validation; **Kavita Yadav:** Methodology, Data curation; **Anupama Kumari:** Methodology, Data curation; **Deepanshu Maheshwari:** Methodology, Data curation; **Pragyan Parimita Rath:** Methodology and resources; **Jaswinder Singh Maras:** Resources, Data curation, Formal analysis; **Asif Mohmmed:** Methodology and resources; **Anupam Kumar:** Resources, Data curation; **Shiv Kumar Sarin:** Conceptualization, Writing - review & editing; **Shvetank Sharma:** Conceptualization, Methodology, Investigation, Resources, Supervision, Writing - review & editing. All the authors have read and approved the final manuscript.

