## Additional file 1 for "Diet-conditioned microbiota enhances fecal microbiota transplantation efficacy in alcoholic liver disease through caproic acid-PPARα signaling"

*Dr. Shvetank Sharma, PhD*

*Associate Professor,*

*Department of Molecular and Cellular Medicine,*

*Institute of Liver and Biliary Sciences, New Delhi-110070, India*

**

***# Equal contribution***

**Contents**

S1: Material and methods

S. Fig1. General characteristics of FMT treated mice

S. Fig2: Histopathology of ileum tissues using Hematoxylin and Eosin (HE) stain

S. Fig3: Gut micorbiota composition profiles

S. Fig4: Hepatic proteomics profiles of FMT treated groups

S. Fig5: General characteristics PPARα inhibitor treated mice

S. Fig6: Comparison of gut microbiota composition profiles in veg-FMT with or without PPARα inhibition

S. Fig7: Veg-FMT increases Caproic acid in the stool and systemic circulation.

S. Fig8: Integrated correlation analysis of microbiota, caproic acid, and host phenotypes.

S. Fig9: Caproic acid supplementation alleviated ethanol-induced hepatic inflammation and lipid accumulation in Huh7 cell line.

S. Fig 10: Biolayer interferometry analysis of the interaction between caproic acid and the PPARα ligand-binding domain.

S. Fig11: General characteristics of Caproic acid with or without PPARα inhibitor treated mice.

**S1: Material and methods**

**S1.1 Triglyceride estimation**

Triglyceride (TG) levels in liver tissue and Huh7 cells were quantified using a colorimetric assay kit based on the glycerol phosphate oxidase–peroxidase (GPO-PAP) method (Elabscience, Cat. No. E-BC-K261-S). For liver samples, ~20 mg tissue was washed with cold normal saline and homogenized in 180 µL anhydrous ethanol at 4 °C, followed by centrifugation at 10,000 × g for 10 min at 4 °C. For cellular TG estimation, Huh7 cells were washed twice with cold PBS, harvested by scraping, homogenized in anhydrous ethanol, and centrifuged under identical conditions.

The resulting supernatants were used for TG estimation. Briefly, 10 µL of sample or standard was mixed with 1000 µL of enzyme working solution and incubated at 37 °C for 10 min. Absorbance was measured at 510 nm using a spectrophotometer. Triglyceride concentrations were calculated using the kit standard and normalized to wet tissue weight (µmol/g) for liver samples or total protein content (mmol/g protein) for cell samples, in accordance with the manufacturer’s instructions.

**S1.2 Gene expression analysis**

Hepatic and ileal recovery post-FMT was assessed using tissue specific gene expression analysis. RNA was isolated from the liver and ileum tissues using TRIzol™ Reagent (Cat#15596026, Thermo, USA). This was followed by cDNA synthesis (Cat#AB1453A, Verso cDNA kit, Thermo, USA) and real-time PCR (Applied Biosystems 7300 real-time PCR system, California, USA). Expression of pro-inflammatory (IL-6, TNFα, IL1β, NF-κB, Tgfβ, TLR4, ACTA2), anti-inflammatory markers (IL10), beta-oxidation (PPARα, PPARγ, Cpt1, Acox1, Acadvl, Acadm, Fabp1) and lipogenesis (Srebpc1, Scd1, Fasn) genes were assessed in hepatic tissue. Ileal tissues were assessed for tight junction (claudin3, occludin, Zona occludens), antimicrobial peptides (Reg3β and Reg3ϒ) genes, and mucin layer (muc2). The ΔΔCT method was used to calculate gene expression using 18s gene expression as control. Amplification was initiated with an enzyme activation step at 94°C for 3 min, followed by 35 cycles consisting of a 30s denaturation at 94°C, a 15s annealing at the temperature of 56°C-60°C for 30 s, and a 45s elongation step at 72°C. Primer sequences and qPCR details are provided in Additional file 2, S-table 1.

**S1.3 Gut microbiota analysis**

Gut microbiota composition was analysed using 16S rRNA sequencing, six biological samples per recipient group were included in the microbiome analysis. Fecal DNA was extracted using a SPINeasy DNA Kit (SKU: 116547050, MP Biomedicals) according to the manufacturer’s protocol. A 16S rRNA sequencing library was constructed according to the 16S metagenomics sequencing library preparation protocol (Illumina, USA), targeting the V3-V4 hypervariable regions of the 16S rRNA gene. All the samples were run together for sequencing on Novaseq 2000 sequencing platform. The reads were trimmed to remove PCR primers and adaptors to finally have a sequence length of 300 bases. Sequences with a PHRED score greater than 30 were considered for downstream analysis. Amplicon sequence variants (ASVs) were identified using DADA2 pipeline. Rare ASVs (<0.01 % counts) were removed while keeping minimum sum of reads within all samples to more than 10,000 and with occurrence in at least 50% samples. SILVA database was used taxonomic reference for ASV classification. Statistical and ecological analysis of the meta-taxonomic data was performed as described by Volant, et al (1). To predict microbial functional potential from 16S rRNA sequencing data, phylogenetic investigation of communities by reconstruction of unobserved states (PICRUSt2) was employed. Amplicon sequence variants were normalized for 16S rRNA gene copy number and used to infer metagenome functional content. Predicted gene families were subsequently mapped to MetaCyc metabolic pathways. Differential pathway abundance between experimental groups was analyzed and visualized using volcano and bidirectional bar plots.

**S1.4 Hepatic proteome LC-MS/MS analysis**

To assess the hepatic response to FMT, we compared the total liver proteome between the groups (n=3 per group). Total proteins were isolated from the 200mg liver tissues. 50µg equivalent protein was subjected to reduction, alkylation followed by digestion for 16–20 hr at 37°C using sequencing-grade modified trypsin:proteins (1µg:20µg w/w). The samples were desalted and subjected to LC-MS/MS analysis. The peptides were eluted by a 3–95% gradient of buffer B (aqueous 80% acetonitrile in 0.1% formic acid) for about 75 min on a 25-cm analytical C18 column (3mm x 100A˚). The peptides were ionized by nano-electrospray and subsequent tandem mass spectrometry (MS/MS) on a Q-Exactive TM Plus (Thermo Fisher Scientific, USA). The MS/MS data was analyzed by Proteome Discoverer (version 2.0, Thermo Fisher Scientific, USA).

Differentially expressed proteins (at ±log2 fold change, p<0.05) were searched in Enrichr (2) to determine significantly associated (p<0.05) KEGG-annotated pathways and associated biological processes.

**S1.5 Confirmation of peroxisome proliferator alpha as the link between liver and gut microbiota**

Outcome of PPARα inhibition under different treatment conditions were tested to confirm its involvement in response to FMT. Ten ALD mice were intraperitonially administered PPARα inhibitor GW6471 (Cat#HY-15372; 10mg/kg body weight, alternate days for two weeks) from the start of 12^th^ week. At the end of 12th week, FMT was performed along with the inhibitor till the end of 13^th^ week. The control group (n=6) received FMT but without inhibitor (Fig. 6A).

For caproic acid validation, total 16 mice were included. In the first group, 8 ALD mice received caproic acid supplementation (2.5% w/w; Cat#153745, Sigma-aldrich, USA) via oral gavage on alternate days for one week, starting after 12th week. In the second group, ALD mice (n=8) were intraperitoneally administered PPARα inhibitor alternatively for two weeks as described above. Caproic acid supplementation was added to the inhibitor treatment at the end of 12th week and continued till the end of 13th week (Fig. 9A).

**S1.6 Western Blot**

Western blot was performed for PPARα to validate the findings of proteomics and also to check the tight junction protein (Zo-1) expressionTotal proteins from the frozen liver and ileum tissue samples (30 mg) were extracted with 300 µl RIPA lysis buffer (Cat#786-490, Gbiosciences, USA) in the presence of a protease inhibitor cocktail (Cat#P8340, Sigma-Aldrich, USA). Proteins in SDS-loading buffer were subjected to SDS-polyacrylamide gel (10%) electrophoresis and transferred to a PVDF membrane (Cat#162-0177, Bio-Rad, USA). Antibodies against PPARα (1:1000; Cat#ITT05301, Immunotag, USA) and beta-actin (1:2000; Cat#sc-47778, Santa Cruz Biotechnology, USA) were used for western blot analysis. The membranes were probed with primary antibodies in 10 ml of blocking buffer (5% skimmed milk) overnight at 4 °C. After washing, the membranes were further probed with the horseradish peroxidase-conjugated secondary antibody (Cat#sc-2357, Santa Cruz Biotechnology, USA) in 10 ml of blocking buffer for 1 h at room temperature. Clarity Western ECL substrate (Cat#170-5060, Bio-rad, USA) was used for chemiluminescence development.

**S1.7 Plasma and stool metabolome**

To find the link between FMT and PPARα activation, we performed plasma and stool metabolome. 50mg stool samples (n=6 per group) were processed using 400µl RIPA lysis buffer (Cat#89901, Thermofisher Scientific) and sonication (30 amplitude; 10 sec on/off pulses in a Helix Biosciences Sonicator, India). Stool lysate and plasma samples were centrifuged at 14,000 rpm for 15 minutes. The supernatants were collected in separate 1.5 mL Eppendorf tubes. In 100 µL of stool lysate and plasma sample, 400 µL of ice-cold methanol was added respectively and kept at -20°C overnight. Samples were then centrifuged at 14,000 rpm for 30 minutes. The supernatant was transferred to new tubes and dried in Speed-Vac at 45°C. The dried supernatant residues were reconstituted in 95% water: 5% internal standards:5% acetonitrile) and then subjected to reverse-phase chromatography in C18 column (Cat#25003102130: 3µm x 2.1mm x 100mm, Thermo Scientific) using ultra-high-performance liquid chromatographic system followed by high-resolution orbitrap mass spectrometry (HRMS, Q Exactive Orbitrap Mass Spectrometer (Thermo Scientific, USA). Metabolites were annotated using Compound Discoverer 3.0 (Thermo Fisher Scientific) and filtered by MAD/median for quality control. Data normalization and scaling were performed in MetaboAnalyst 5.0 (http://metaboanalyst.ca), followed by PLS-DA, heatmap generation, and multivariate statistical analyses to identify significant metabolic features.

**S1.8 In-vitro experiment to confirm the link between caproic acid and PPARα:**

To confirm the role of caproic acid and PPARα in ameliorating ethanol-induced liver injury, Huh7 hepatocellular carcinoma cell line was used. Cells were grown in Dulbecco's Modified Eagle Medium (DMEM) media (Cat#11995-040, Gibco, Thermo Fisher Scientific, USA) supplemented with 10% Fetal Bovine Serum (FBS, Cat#26140079, Gibco, Thermo Fisher Scientific, USA) and maintained at 37 °C in a humidified incubator under 5% carbon dioxide atmosphere.

To assess cell viability, approximately 2x10^4^ cells/mL were seeded into 96-well tissue culture plates and incubated at 37°C for 24 h prior to treatment. After 24hrs, spent media was removed and fresh media with different concentrations of ethanol (Cat#MB228, HIMEDIA, India) (20mM, 50mM, 80mM, 100mM, 150mM, 200mM) or Caproic acid (0.5mM, 900µM, 2.2mM, 4.5mM) was added into various wells. Untreated wells served as control. After the incubation at 37°C of 72hrs to the ethanol treated and 24 hrs to the caproic acid treated, 10µl of MTT reagent (Cat#3361, SRL, India, working conc. 5mg/ml) were added into each well and further incubated for 3hrs at 37°C. The supernatant was removed and the purple formazan crystals dissolved in 100 µL of dimethyl sulfoxide (Cat#D2650, Sigma-aldrich, USA). The optical density of the solution was measured at 570 nm using a micro-plate reader. Treatment readings were normalized against blank wells (without cells). The cell viability of treated samples was expressed as a percentage relative to untreated control samples. All experiments were conducted in triplicate.

The experimental groups were cultured in high-glucose DMEM with 10% FBS under varying conditions as follows. The control group was maintained in standard medium and caproic acid (CA; Cat#153745, Sigma-aldrich, USA) control group received 900µM CA for 24 hours. The treatment groups were exposed to 100 mM ethanol for 72 hours, and further divided into different groups. The abstinence group was exposed to ethanol-free medium for next 24 hours. The Ethanol+CA group was exposed to 900µM CA supplemented with ethanol for next 24 hours. The Abst+CA group exposed to CA supplementation alone (900µM) for 24 hours. Lastly, the Abst+(PPARα inhibitor)+CA group exposed to PPARα inhibitor (10µM) with CA supplementation (900µM) for 24 hrs.

**S1.9 Oil Red O staining:**

Since, caproic acid and PPARα are involved in fatty acid β-oxidation, we conducted oil red O staining. The Huh7 cells were sub-cultured with 10^4^ cells per well in 24-well culture plate following 24 hrs incubation in the 37 °C incubator containing 5% CO_2_. Wells were subsequently divided into six groups as the blank control group, caproic acid alone group, ethanol group, Abstinence group, EtOH+CA group, Abst+CA group and Abst+PPARα inhibitor+CA group.

After incubation, the culture medium was removed, and the experiment was conducted as follows: Cells were washed using 0.5ml PBS three times; 500ul of 4% paraformaldehyde was added into each well for a 3hrs fixation and then washed with 0.5ml ddH_2_O three times. Each well received 500µl of 60% isopropanol and was then incubated for 5 min at room temperature. Isopropanol was decanted and 500µl Oil red O working solution was added into each well and left for staining for 15 min at room temperature. Wells were cleaned with five rinses of ddH_2_O. Cell morphology and the distribution of lipid droplets were observed with the inverted microscope. Lipid accumulation was quantified by washing the cells thrice with 60% Isopropanol. Oil Red O staining was extracted by treating the cells with 100% isopropanol for 5 min. with gentle rocking. Absorbance of each well was determined at 492 nm.

**S1.10 RNA extraction and qPCR analysis:**

To investigate the inflammatory, fatty acid β-oxidation and lipogenesis, we performed gene expression analysis in these cells. Cells were collected at defined time points and total RNA was isolated using Trizol manual method. Reverse transcription was performed 4 µg total mRNA in a final volume of 20 µl reaction using RevertAid First Strand cDNA Synthesis Kit (#K1622,Thermo Fisher Scientific, USA). Real-time polymerase chain reaction (PCR) as previously described was performed for markers IL-6, TNFα, IL1β, NF-κB, IL10, PPARα, Cpt1, Acox1, Fabp1, Srebpc1, Scd1, Fasn, PLIN2, APOB and MTTP using ready-to-use SYBR green according to the manufacturer’s instructions. The relative expression levels compared with control cells (relative expression value set to one) were defined by the 2-ΔΔCt method. Hypoxanthine guanine phosphoribosyltransferase (HPRT) was used as housekeeping gene. Primer sequences are provided in Additional file 2, S-table 1.

**S1.11 XF Mito Fuel Flex Test:**

Since we found increase in PPARα and caproic acid and both participate in mitochondrial fatty acid β-oxidation, we performed a mitochondrial fuel flex test. It was conducted to evaluate the cells' dependency on fatty acid oxidation. The assay was performed following the instructions provided by Seahorse Bioscience (Agilent, USA) which is used to study the cellular bioenergetics. Previously described cell line treatment groups were sequentially treated with etomoxir (4 μM), an inhibitor of fatty acid oxidation, and UK5099 (2 μM), an inhibitor of pyruvate transport, while the oxygen consumption rate (OCR) was measured before and after the addition of each compound. Data analysis was performed using XF Wave software version 2.6 (Agilent, USA). Cellular protein content was quantified using the BCA assay kit (Cat#500-0207, Bio- Rad Laboratories, USA) following standard protocols, and all OCR data were normalized to protein levels.

S1.12 Biolayer Interferometry (BLI)

In order to gauge direct interaction between PPARα and Caproic acid, we have used Bio-layer interferometry (BLI) (Octet®R8 system, Sartorius). 500µg of His-tagged Human-PPARα (MedChemExpress HY-P7996) was immobilized onto a Ni-NTA chip for 600s and the unbound protein was washed using 1X Phosphate Buffered Saline buffer. We prepared all the different concentrations of Caproic acid in 1X PBS for the titrations. 31.25µM, 62.5µM, 125µM, 250µM, and 500µM concentrations were measured for binding.

All the concentrations for each compound were prepared and loaded in the Greiner 96-well plate black F-bottom plate (Cat. No. 655077). The kinetics of the interaction was monitored for 180s each for the association and the dissociation curves by dipping the sensors in a series of increasing concentrations of Caproic acid. 1X PBS was used as the dissociation buffer. A control sensor with the immobilized protein was used in parallel and was used to normalize the data. Washing in 350 mM EDTA regenerated the sensors for the next cycle of binding kinetics. The experiments were performed at 25°C. All the real-time data was analyzed and KD values were determined using the Octet® Evaluation software using 1:1 fitting curves.**S1.13 Statistical Analyses**

Graph Pad Prism version 8 was used for statistical analyses. Continuous variables were compared between two groups using either the Student's t-test or the Mann-Whitney test, and among more than three groups using ANOVA or the Kruskal-Wallis test, as appropriate. The categorical variables were tested using either the Pearson chi-square test or the Fisher exact test, as appropriate. Spearman correlation (r^2^ > 0.7, p < .05) was plotted using Corr plot in R (version 4.4.0). For evaluating 16s data, one-way ANOVA was used for continuous variables, and the chi-square test was used for categorical variables. Bray-Curtis distances were used for the principal coordinate analysis (PCoA). p<0.05 was taken as a significant statistical measure.

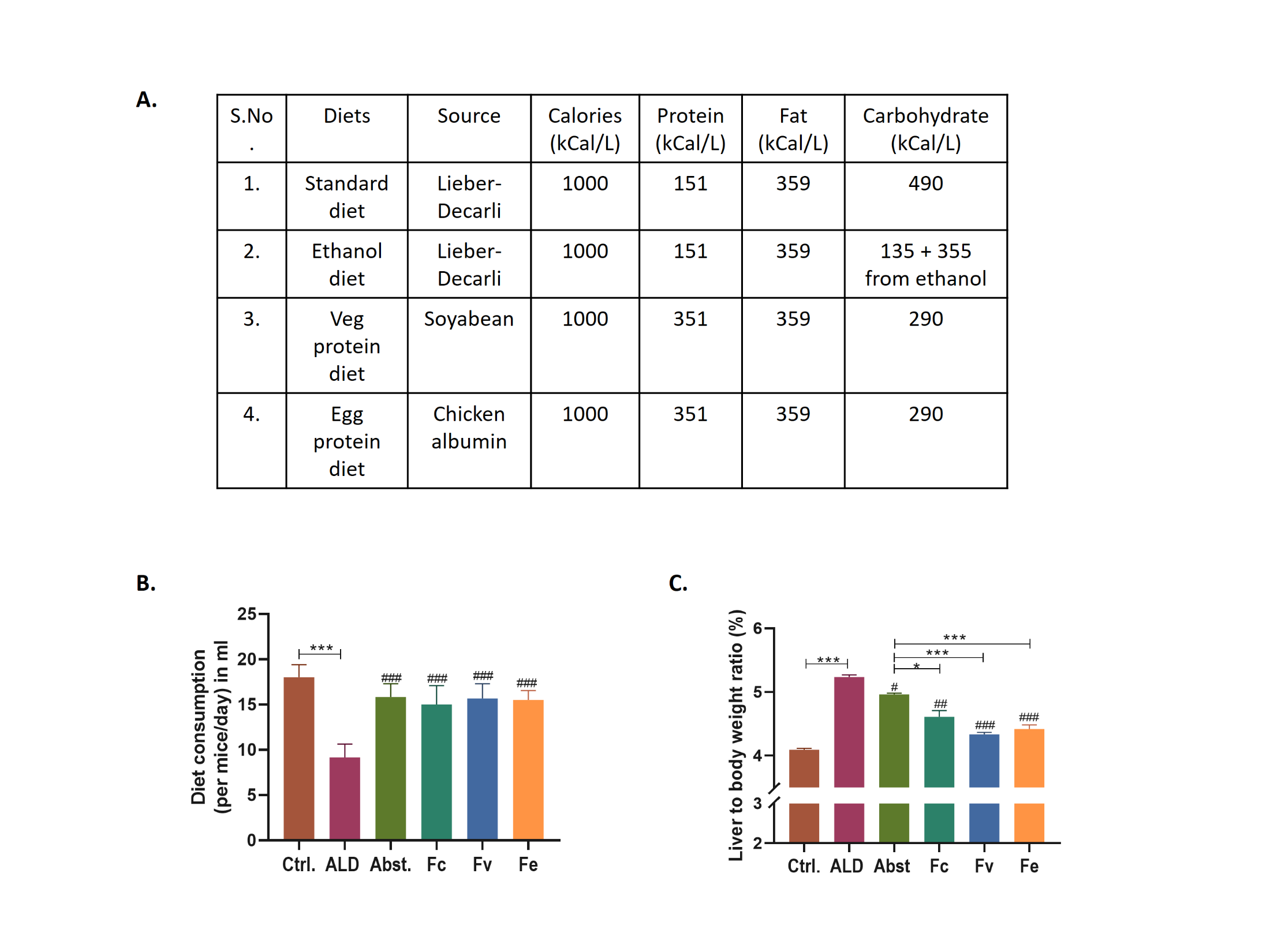

**S. Fig 1: General characteristics of FMT-treated mice.** **A)** Calorie profile of diets. **B)** Diet consumption per mice per day remains unchanged irrespective of the treatment. **C)** Liver to body weight ratio reduced post protein-FMT (liver index).

Data are presented as mean ± SEM (standard error mean). *p < 0.05, **p < 0.01, ***p < 0.001 shows inter-group statistics; #p < 0.05, ##p < 0.01, ###p < 0.001 with respect to the ALD group(one-way ANOVA followed by Tukey’s multiple comparison test).

*ALD: Alcohol-related liver disease, Abst: abstinence, Fc: Std-FMT, Fe: Egg-FMT, Fv: Veg-FMT*

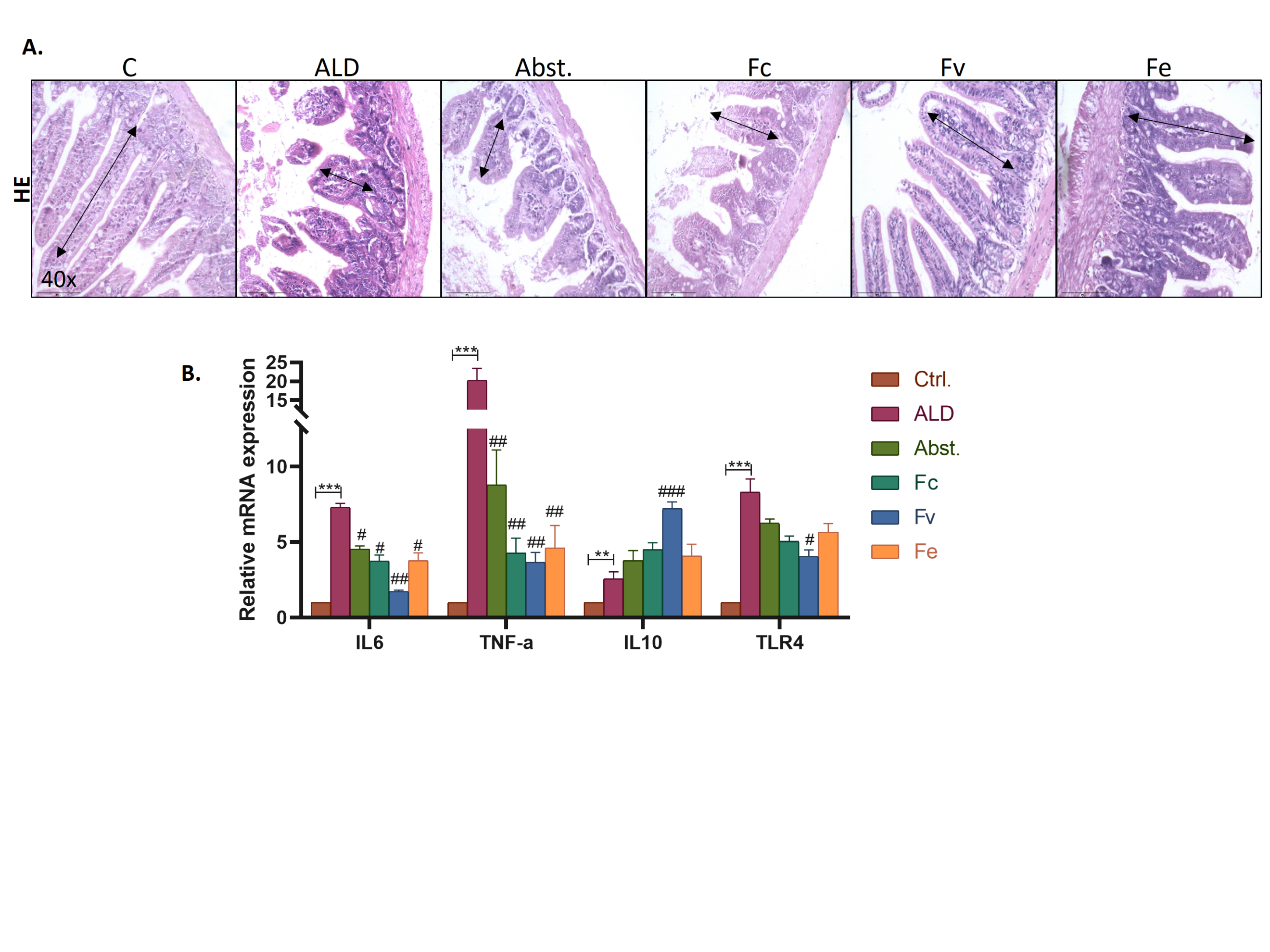

**S. Fig 2: (A)** Histopathology of ileum tissues using Hematoxylin and Eosin (HE) stain. Arrow shows the reduced villus length in ALD group which was increased after protein-FMT. **(B)** mRNA expressions of pro-inflammatory reduced and anti-inflammatory increased after veg-protein FMT.

Data are presented as mean ± SEM (standard error mean). *p < 0.05, **p < 0.01, ***p < 0.001 shows inter-group statistics; #p < 0.05, ##p < 0.01, ###p < 0.001 with respect to the ALD group(one-way ANOVA followed by Tukey’s multiple comparison test).

*IL: Interlukin; TNF-α: tumor necrosis factor alpha; TLR 4: Toll like receptor 4.*

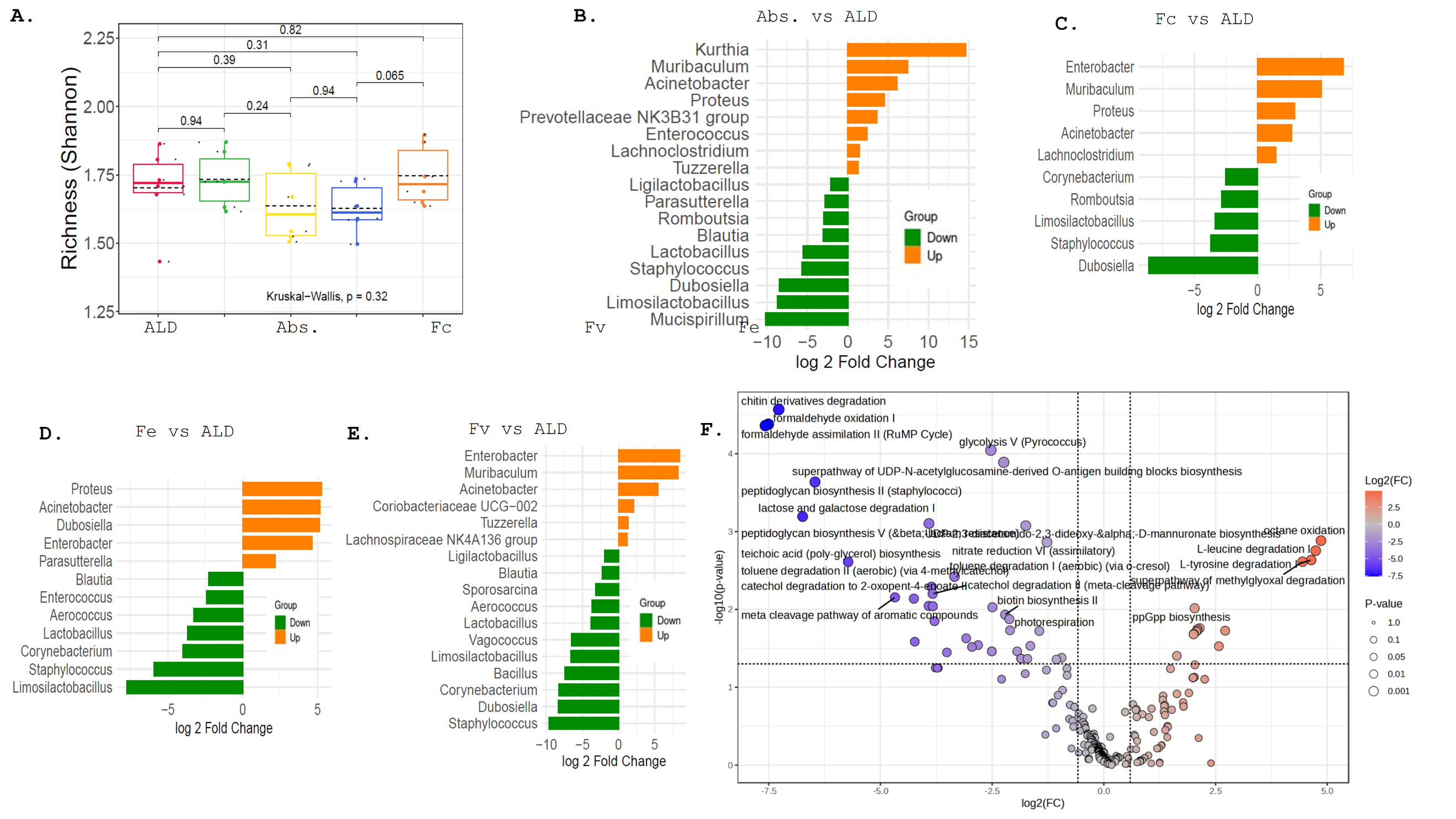

**S. Fig 3: Gut micorbiota composition profiles.** **A)** Alpha diversity using Shannon index showing non-significant changes in the treatment groups. **B)** Differential gut bacteria (FC>2, p<0.05) between Abst and ALD **C)** Fc vs ALD **D)** Fe vs ALD **E)** Fv vs ALD. **F)**. Volcano plot of predicted microbial metabolic pathways in veg-FMT.

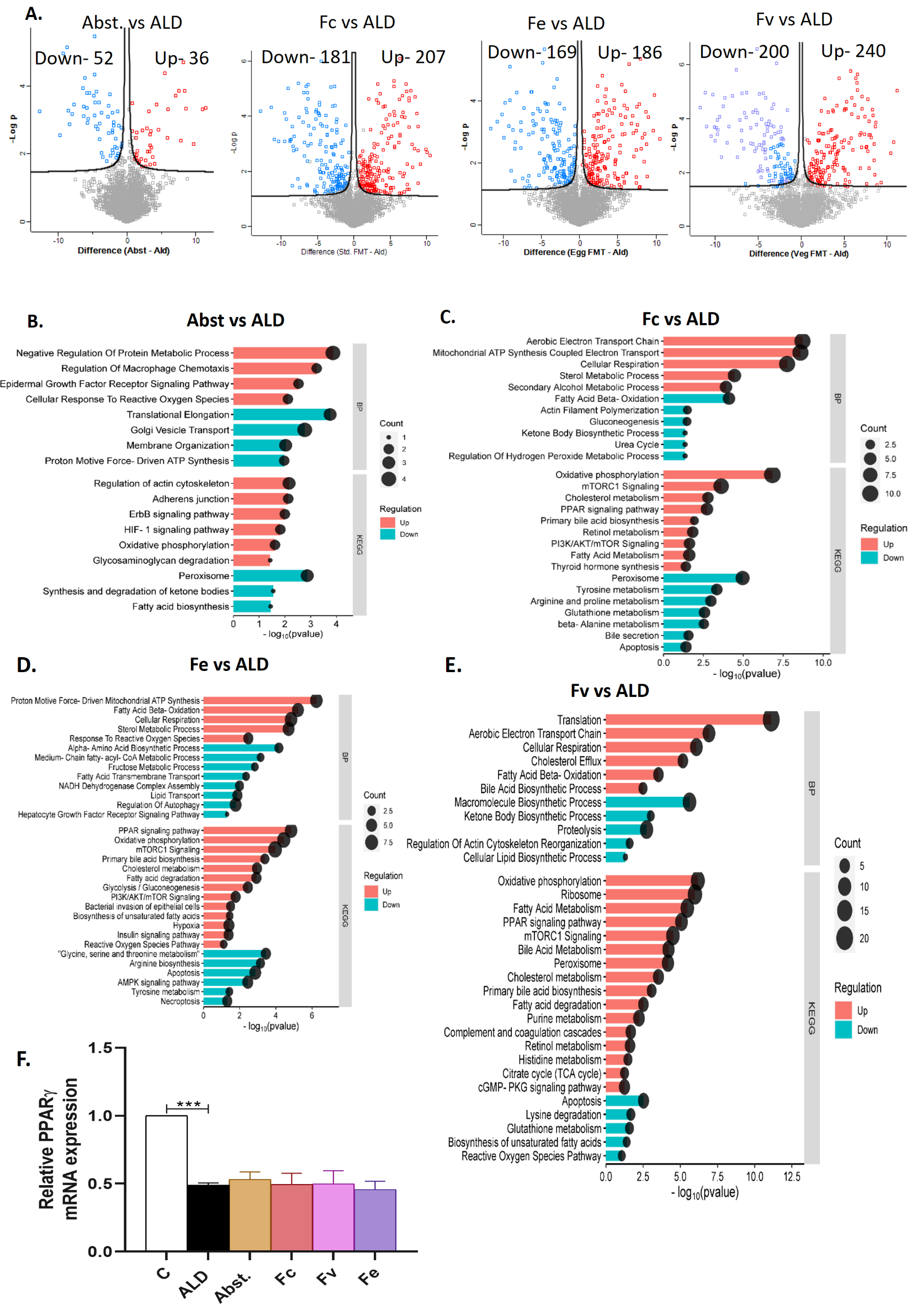

**S. Fig 4: Hepatic proteomics profiles of FMT treated groups.** **A)** Differential proteins (FC>2, p<0.05) between Abst and ALD, Fc vs ALD, Fe vs ALD and Fv vs ALD, **B)** Differential pathways (FC>2, p<0.05) between Abst and ALD. **C)** Fc vs ALD **D)** Fe vs ALD **E)** Fv vs ALD **F)** mRNA expression of PPARϒ

Data are presented as mean ± SEM (standard error mean). *p < 0.05, **p < 0.01, ***p < 0.001 shows inter-group statistics

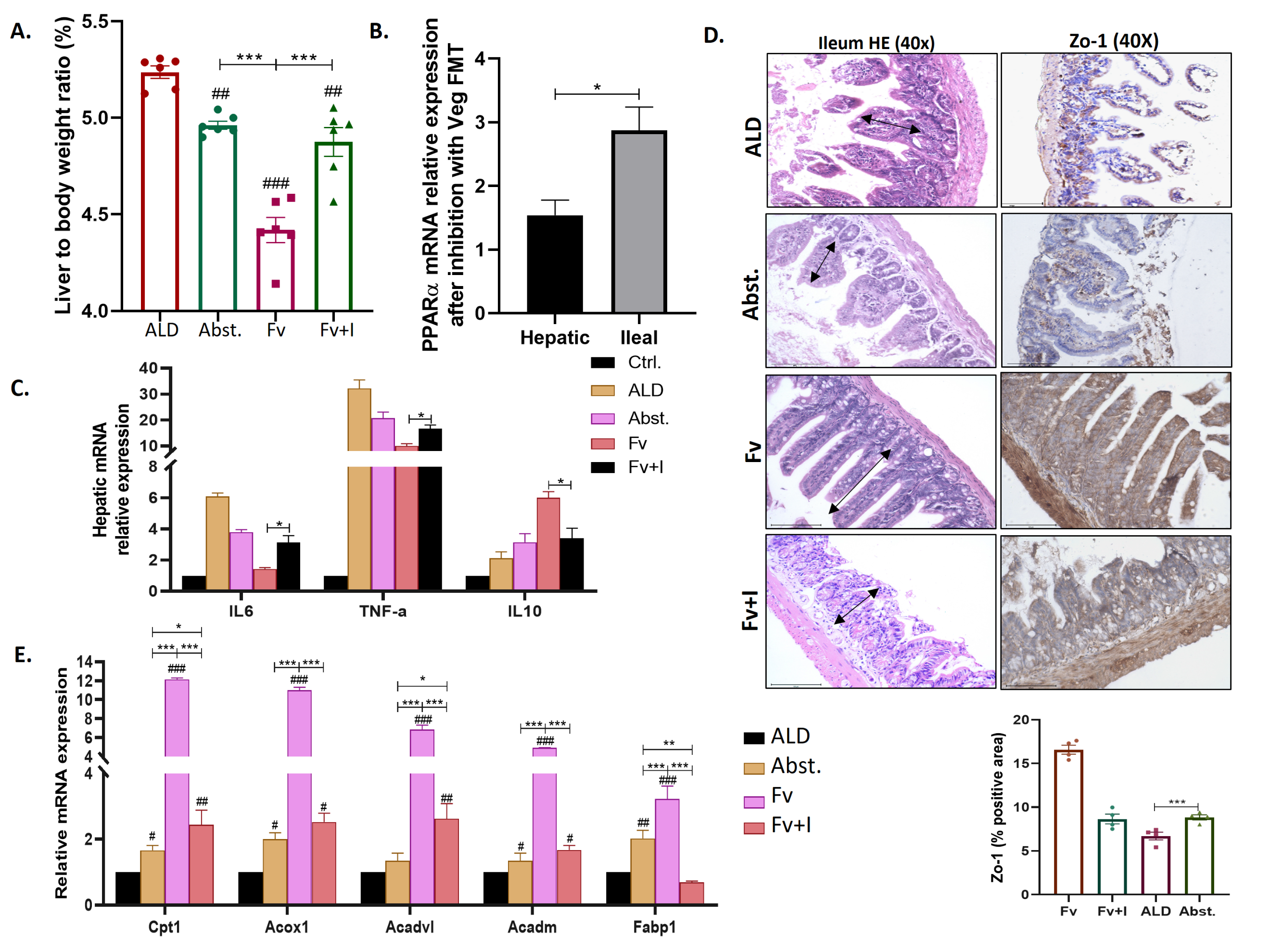

**S. Fig 5: General characteristics PPARα inhibitor treated mice** **A)** Liver to body weight ratio reduced significantly post veg-FMT. **B)** mRNA expression of hepatic PPARα significantly reduced in the presence of PPARα inhibitor after veg-FMT compared to ileal PPARα expressions. **C)** Histopathology of ileum tissues using Hematoxylin and Eosin (HE) stain. Arrow shows the reduced villus length in ALD group which was increased after protein-FMT and these changes were reversed after PPARα inhibitor and **D)** Zo-1 protein expression by immunohistochemistry showing improved barrier integrity in protein-educated FMT and these imprvements were not shown after pparα inhibition with statistical analysis. **E)** mRNA expressions of pro-inflammatory reduced and anti-inflammatory increased after veg-protein FMT and these changes were not observed after PPARα inhibiton.

**F)** mRNA expression of PPARα target genes were significantly increased in veg-FMT and this effect of veg-FMT was reduced in the presence of PPARα inhibitor.

Data are presented as mean ± SEM (standard error mean). *p < 0.05, **p < 0.01, ***p < 0.001 shows inter-group statistics; #p < 0.05, ##p < 0.01, ###p < 0.001 with respect to the ALD group(one-way ANOVA followed by Tukey’s multiple comparison test).

*Cpt1: Carnitine palmitoyltransferase 1, Acox1: Acyl-Coenzyme A oxidase 1, palmitoyl, Acadvl: Acyl-Coenzyme A dehydrogenase, very long chain, Acadm: Acyl-Coenzyme A dehydrogenase, medium chain, Fabp1: Fatty acid binding protein 1, IL: Interlukin; TNF-α: tumor necrosis factor alpha; TLR 4: Toll like receptor 4.*

this

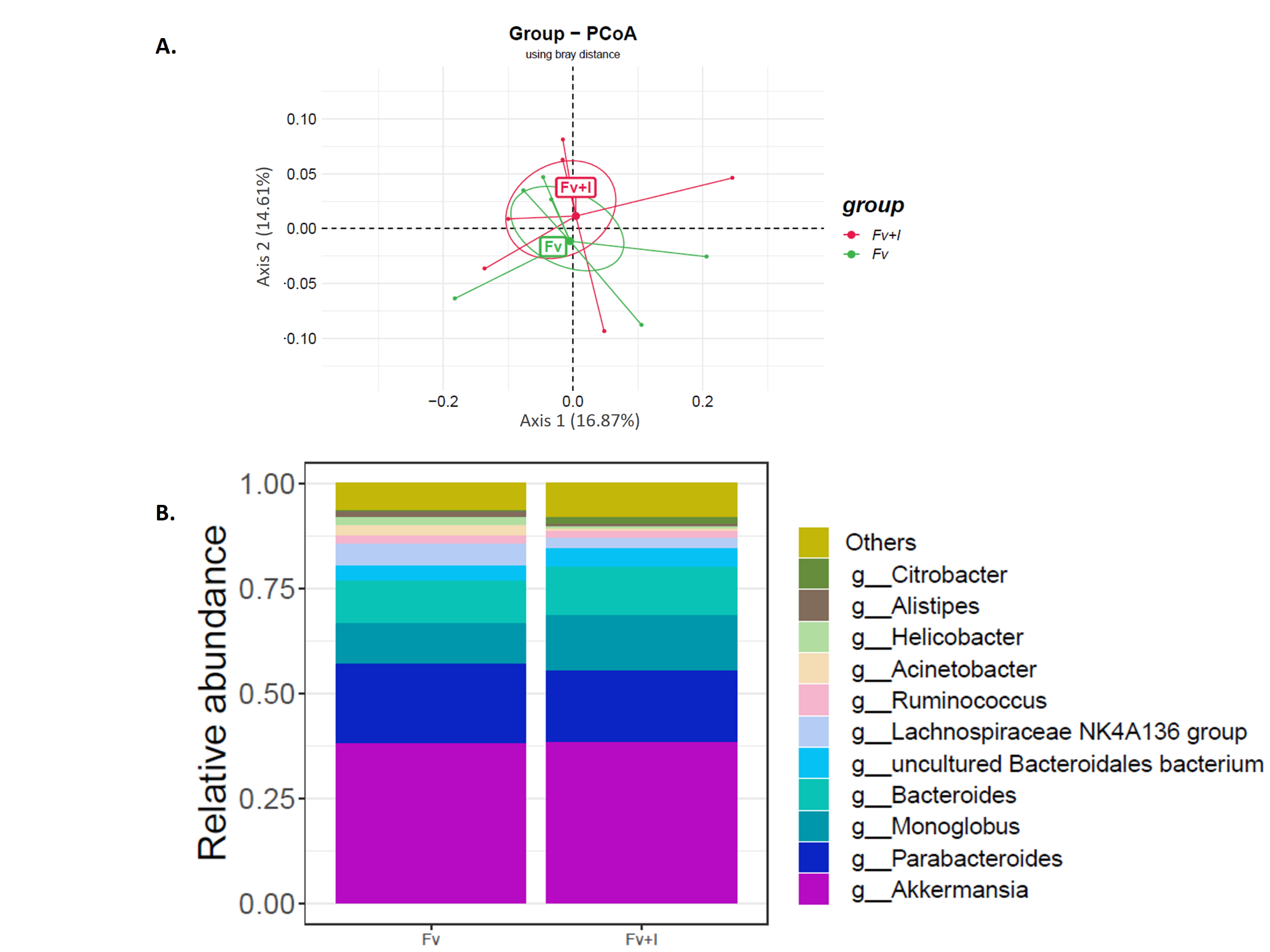

**S. Fig 6: Comparison of gut microbiota composition profiles in veg-FMT with or without PPARα inhibition.** **A)** Beta diversity using bray-Curtis distance showing no significant (p=0.6) variance between the groups. **B)** Relative comparison showing almost similar abundance of bacterial genera post veg-FMT in both with or without PPARα inhibitor groups.

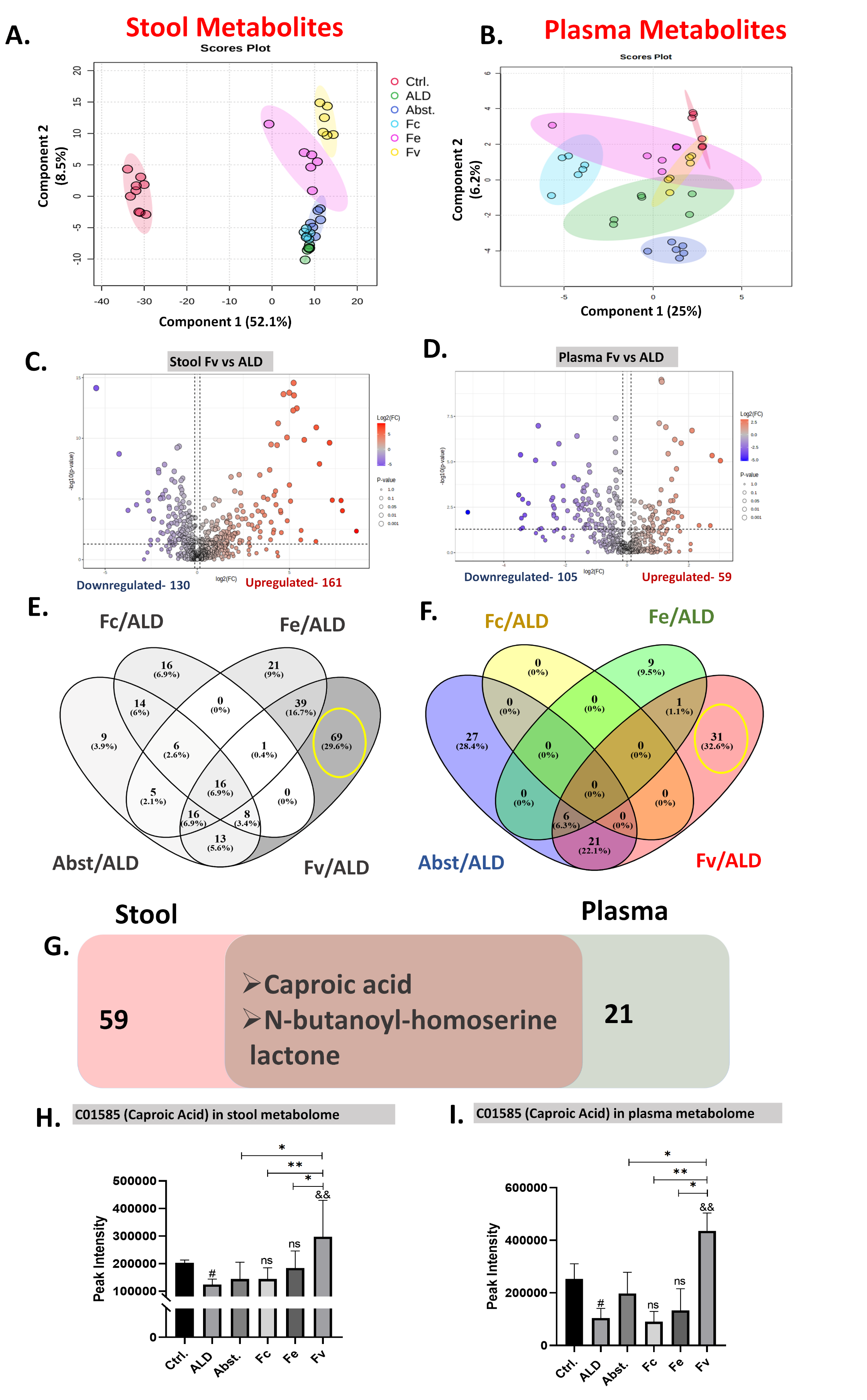

**S. Fig. 7:** **Veg-FMT increases Caproic acid in the stool and systemic circulation.** PCoA of **(A)** stool and **(B)** plasma metabolites showing 60% variance in stool and 31.2% variance in plasma metabolome. Differential metabolites showing significantly (FC>1.1, p<0.05) upregulated and downregulated metabolites in (**C)** stool and (**D)** plasma. Venn diagram showing unique and common metabolites among different FMTs and abst. in **(E)** stool and **(F)** plasma, with 69 stool and 31 plasma metabolites exclusively and significantly increased (FC>1.5, p<0.05) by Fv. **(G)** 2 commonly upregulated metabolites were found between stool and plasma of Fv. Peak intensity of caproic acid showed higher expression in Fv both in **(H)** stool and **(I)** plasma

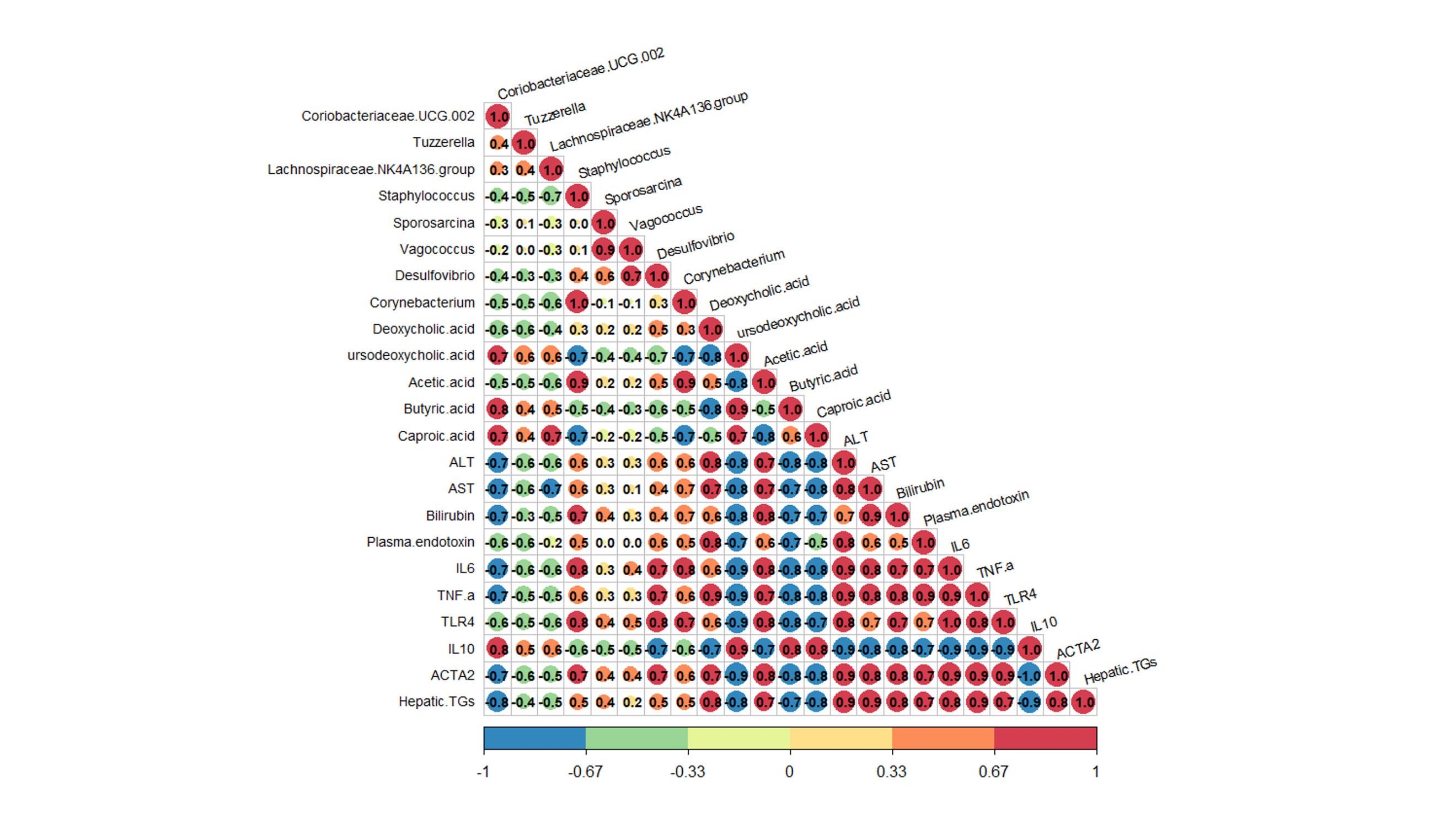

**S. Fig 8: Integrated correlation analysis of microbiota, caproic acid, and host phenotypes.** Spearman correlation analysis was performed using microbiome, metabolomic, and phenotypic data. The heatmap shows correlations between selected bacterial taxa, caproic acid abundance, and hepatic injury, inflammatory, and metabolic parameters. Color intensity represents the strength of the correlation coefficient (ρ), with positive and negative associations indicated by the direction of the correlation. Significant associations were identified using FDR-adjusted q < 0.05.

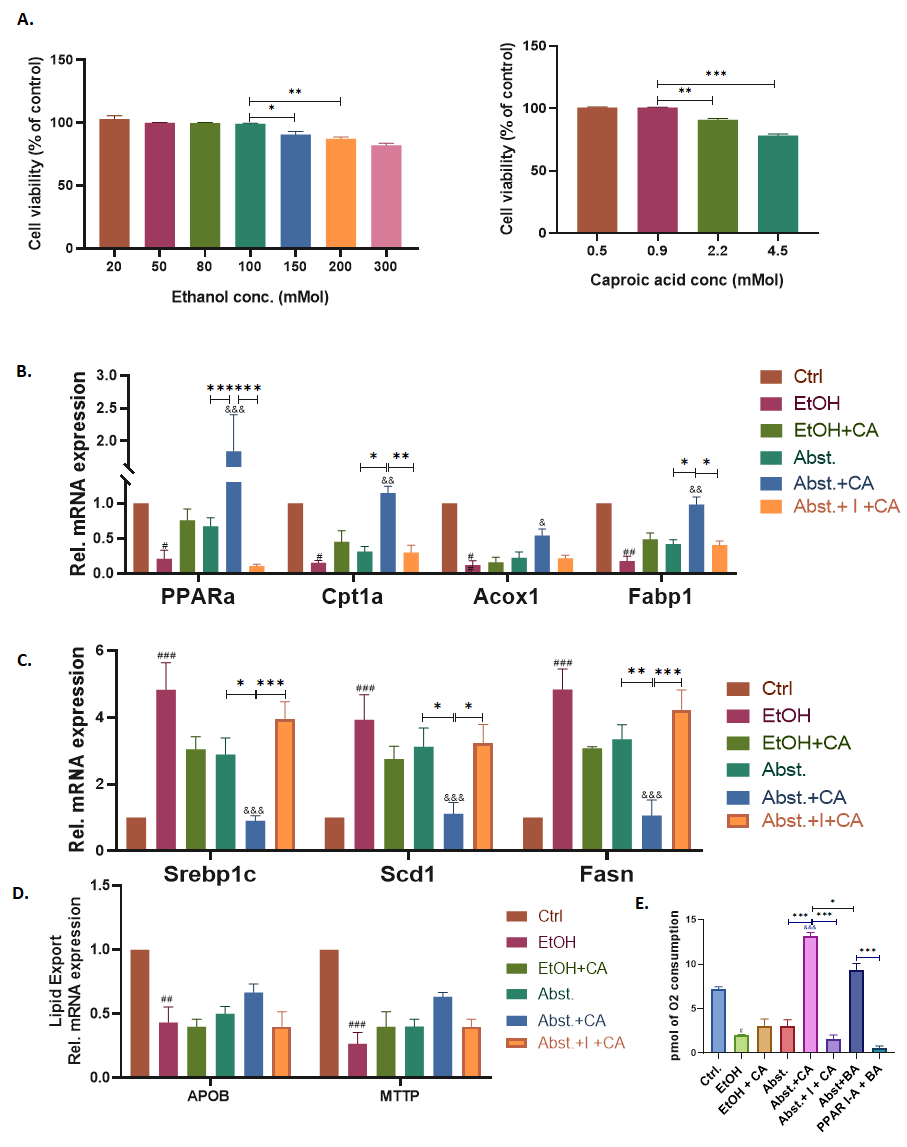

**S. Fig 9:** **Caproic acid supplementation alleviated ethanol-induced hepatic inflammation and lipid accumulation in Huh7 cell line. A)** Cell viability assay on different concentrations of ethanol and caproic acid showing less toxicity effects on 100mM and 0.9mM concentrations, respectively. **(B)** The mRNA expression of PPARα target genes involved in fatty acid β-oxidation was increased, and **(C)** lipogenesis was decreased in CA-supplemented cells, but these effects were reversed in the presence of a PPARα inhibitor (I). **D)** mRNA expressions of lipid export genes (ApoB and MTTP) were decresed after ethanol treatment. **(E)** Mitofuel flex showed, CA was more dependent on fatty acid β-oxidation then butyric acid (BA). Data are presented as mean ± SEM (standard error mean). *p < 0.05, **p < 0.01, ***p < 0.001 shows inter-group statistics; #p < 0.05, ##p < 0.01, ###p < 0.001 with respect to the control. &p < 0.05, &&p < 0.01, &&&p < 0.001 with respect to the ethanol treatment (one-way ANOVA followed by Tukey’s multiple comparison test). *Cpt1: Carnitine palmitoyltransferase 1, Acox1: Acyl-Coenzyme A oxidase 1, Fabp1: Fatty acid binding protein 1, PPARα: peroxisome proliferator-activated receptor alpha, Scd1: stearoyl-coenzyme A desaturase 1, Srebp1c: Sterol Regulatory Element Binding Protein 1c, Fasn: fatty acid synthase, APOB: Apolipoprotein B,MTTP:* *Microsomal Triglyceride Transfer Protein.*

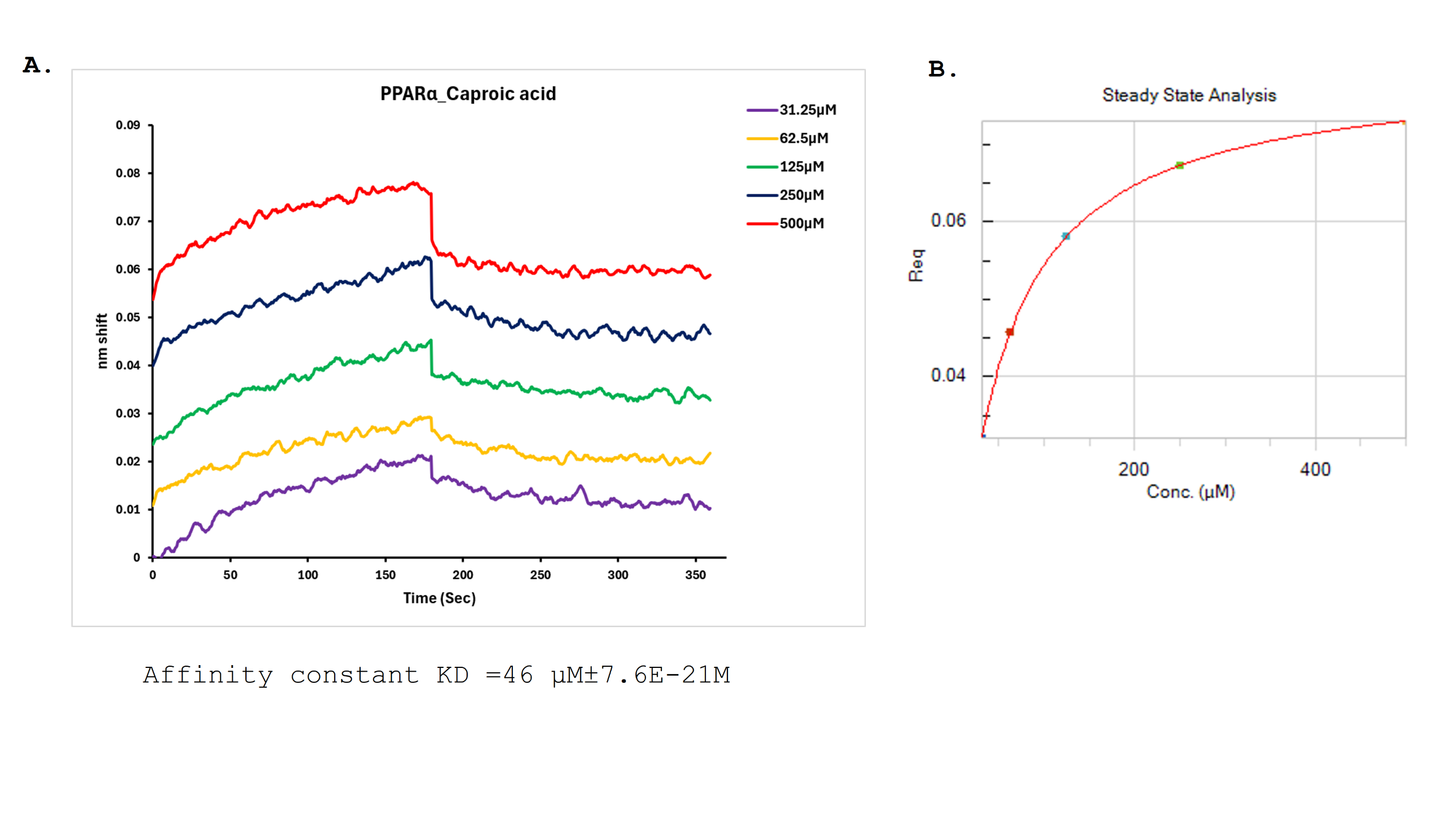

**S. Fig 10: Biolayer interferometry analysis of the interaction between caproic acid and the PPARα ligand-binding domain. (A)** BLI sensorgrams showing concentration-dependent binding responses of recombinant PPARα ligand-binding domain (PPARα-LBD) to increasing concentrations of caproic acid (31.25, 62.5, 125, 250, and 500 µM). The association phase was followed by a dissociation phase in buffer. **(B)** Steady-state analysis showing a concentration-dependent, saturable binding response of caproic acid to PPARα-LBD. The fitted apparent KD was approximately 46 µM.

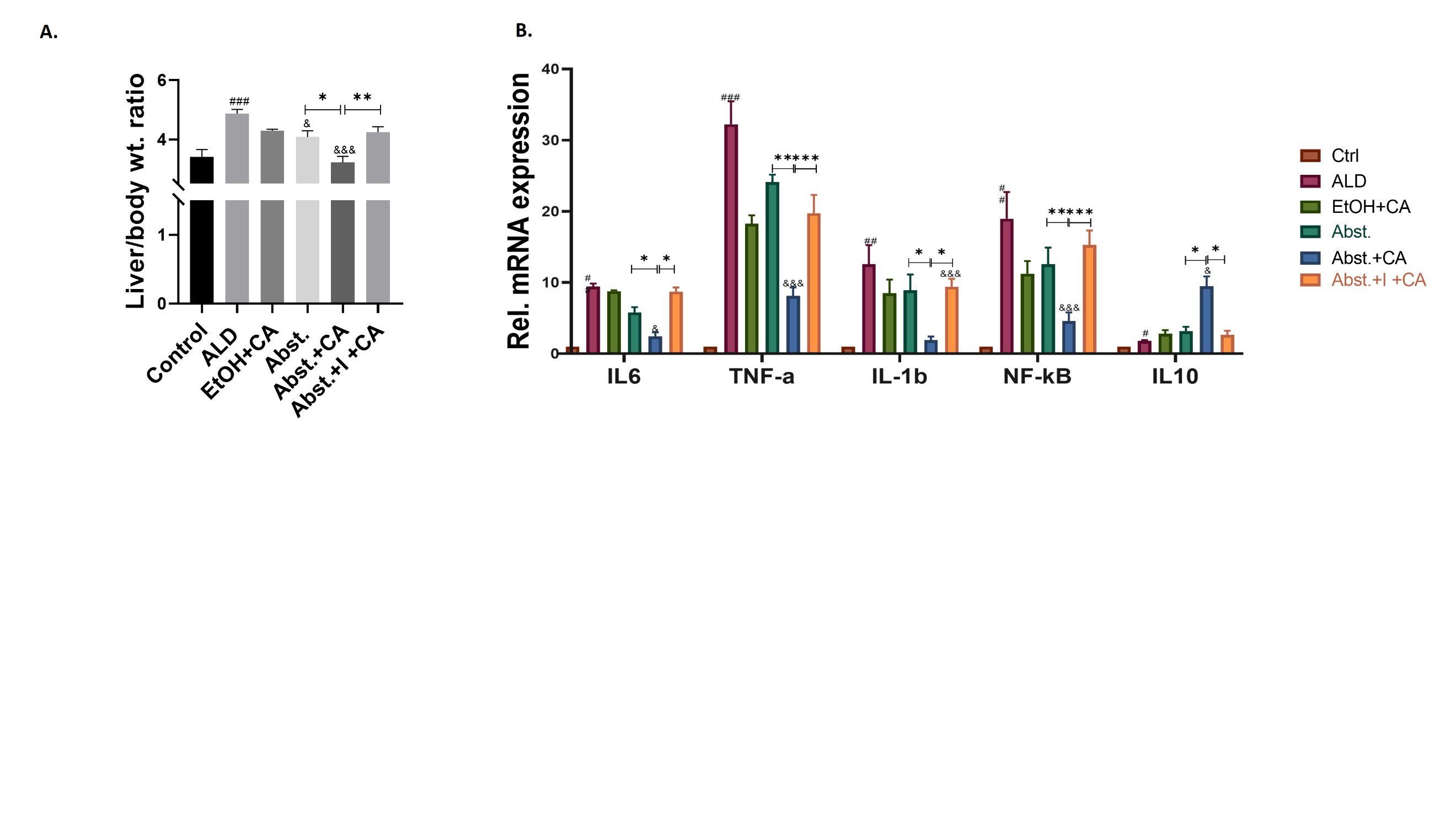

**S. Fig 11: General characteristics of Caproic acid with or without PPARα inhibitor treated mice** **A)** Liver to body weight ratio showed reduction after CA supplementation **B)** Hepatic mRNA expressions of pro-inflammatory markers reduced and anti-inflammatory marker increased in CA group. These changes were found to be reversed in the presence of PPARα inhibitor. Data are presented as mean ± SEM (standard error mean). *p < 0.05, **p < 0.01, ***p < 0.001 shows inter-group statistics; #p < 0.05, ##p < 0.01, ###p < 0.001 with respect to the control. &p < 0.05, &&p < 0.01, &&&p < 0.001 with respect to the ethanol treatment (one-way ANOVA followed by Tukey’s multiple comparison test).

*IL: Interlukin; TNF-α: tumor necrosis factor alpha; NF-kB: Nuclear factor kappa-light-chain-enhancer of activated B cells.*
